# Towards In-Silico CLIP-seq: Predicting Protein-RNA Interaction via Sequence-to-Signal Learning

**DOI:** 10.1101/2022.09.16.508290

**Authors:** Marc Horlacher, Nils Wagner, Lambert Moyon, Klara Kuret, Nicolas Goedert, Marco Salvatore, Jernej Ule, Julien Gagneur, Ole Winther, Annalisa Marsico

**Affiliations:** Computational Health Center, Helmholtz Center Munich, Germany; Department of Biology, University of Copenhagen, Denmark; Department of Informatics, Technical University of Munich, Garching, Germany; National Institute of Chemistry, Ljubljana, Slovenia; The Francis Crick Institute, London, UK; Helmholtz Association - Munich School for Data Science (MUDS), Munich, Germany

**Keywords:** CLIP-seq, Deep Learning, Computational Biology, Protein-RNA Interaction

## Abstract

Unraveling sequence determinants which drive protein-RNA interaction is crucial for studying binding mechanisms and the impact of genomic variants. While CLIP-seq allows for transcriptome-wide profiling of *in vivo* protein-RNA interactions, it is limited to expressed transcripts, requiring computational imputation of missing binding information. Existing classification-based methods predict binding with low resolution and depend on prior labeling of transcriptome regions for training. We present RBPNet, a novel deep learning method, which predicts CLIP crosslink count distribution from RNA sequence at single-nucleotide resolution. By training on up to a million regions, RBPNet achieves high generalization on eCLIP, iCLIP and miCLIP assays, outperforming state-of-the-art classifiers. CLIP-seq suffers from various technical biases, complicating downstream interpretation. RBPNet performs bias correction by modeling the raw signal as a mixture of the protein-specific and background signal. Through model interrogation via Integrated Gradients, RBPNet identifies predictive sub-sequences corresponding to known binding motifs and enables variant-impact scoring via in silico mutagenesis. Together, RBPNet improves inference of protein-RNA interaction, as well as mechanistic interpretation of predictions.

## 1 Background

RNA-binding proteins (RBPs) are a family of proteins that bind to coding and non-coding transcripts, usually through recognition of short sequence or structural features commonly known as motifs [8]. To date, over 2, 000 proteins have been experimentally identified as RNA-binding, rendering it one of the largest cellular protein groups [13]. RBPs are involved in every aspect of post-transcriptional regulation, including modification, stabilization, localization, splicing and translation of RNAs [20]. Misregulation of RBPs, as well as mutations in their amino acid sequence or the sequence of their RNA targets, has been associated with an abundance of human diseases, including neurological and psychiatric disorders [42]. Therefore, uncovering binding preferences and RNA targets of RBPs is crucial for understanding the role of RBPs in post-transcriptional regulatory pathways and for quantifying the impact of their dis-regulation in context of human disease. Nowadays, the most common protein-centric experimental approach to profile RNA-binding *in vivo* is via individual-nucleotide resolution Cross-Linking and Immunoprecipitation followed by sequencing (CLIP-seq) [22, 54] and its derivatives, which enables transcriptome-wide detection of protein-RNA interactions for a protein of interest. Variants of CLIP-seq, including *individual-nucleotide* and *enhanced* CLIP (iCLIP and eCLIP, respectively), further allow for detection of protein-RNA crosslinking events at single-nucleotide resolution. Here, in brief, cells are radiated with UV light, forming covalent cross-links between RNA molecules and bound proteins. Protein/RNA complexes are then purified with protein-specific antibodies and the bound proteins are digested. Subsequently, the bound RNA molecules are reverse-transcribed to cDNA, followed by high-throughput sequencing. Reverse-transcription often truncates at the cross-linked site due to a small peptide remaining bound to the site after protein digestion. After alignment of reads to the reference genome, this leads to an accumulation of read-starts at one nucleotide down-stream of the cross-linked site. The resulting nucleotide-wise count signal can then be used to identify binding sites of the RBP of interest as RNA regions where the signal is significantly higher than expected, given some background model [49]. CLIP-seq data is commonly post-processed with peak callers, which identify a number of regions of enriched signal (also referred to as *binding sites*), usually in the order of thousands. As peak calling is subject to unspecific cross-linking events in the underlying data, inferring target-specific signal from CLIP data is crucial. Multiple studies have identified a number of CLIP-associated biases, including background signal from abundant RNAs that are not properly washed during library preparation, library contamination with bound RNAs of other RBPs and strong UV crosslinking bias towards single-stranded uridine-rich motifs [16]. CLIP-seq experiments are often paired with a control, to account for unspecific background signal. For instance, the eCLIP protocol is designed to generate a size-matched input (SMInput) control by omitting the IP step such that the resulting library represents RNA fragments crosslinked to a mixture of background proteins with a similar molecular weight as the target RBP. Therefore, eCLIP SMInput data is a powerful resource to correct computational methods for experimental bias and reduce the number of detected unspecific binding events for an RBP of interest.

Analysis of experimentally identified RNA binding-sites can give insight into both the functional role of the RBP as well as the RNA sequence and structure feature by which it identifies (and binds to) its RNA targets. In addition, knowledge of the binding preference of an RBP enables imputation of protein-RNA interaction on RNAs not present at the time of the experiment. While traditional methods for *de novo* motif finding [3] or more sophisticated generative models [19] analyze identified RNA binding sites directly, for instance by aggregating enriched sub-sequences into motif position-weight matrices (PWMs), more recent model-based approaches [25, 37] attempt to model RBP-binding as a function of RNA sequence for a given protein of interest. That is, model-based approaches attempt to find a function of the form *f* : *RNA* → *σ*(*CLIP*), where *σ* is some post-processing function on the raw experimental CLIP, for instance a peak caller. Model-based analysis of RBP binding has several key advantages over traditional model-free approaches. First, the availability of a model allows for imputation of missing binding site information. As only a fraction of transcripts may be expressed in the experimental cell type at a given time, CLIP-seq experiments generally draw an incomplete picture of an RBP’s binding landscape. Here, predictions from RBP binding models may aid researchers in generalizing their analysis to unobserved transcripts by providing candidate sites of protein-RNA interaction. The ability to impute missing binding information on arbitrary RNA sequences is especially relevant for the analysis of RBP-binding to foreign sequences, such as foreign RNAs derived from viruses [21]. Second, the model *f* represents a simplified abstraction of the *in vivo* biology, that is, the recognition of a binding site in a target RNA sequence by the protein of interest. Besides identifying drivers of RNA target recognition (e.g. binding motifs), the ability to study *f* enables “what-if” analysis, allowing one to explore how changes in the RNA input sequence affect RBP binding. For instance, researchers may investigate the impact of single-nucleotide variants (SNVs) on RBP binding *in silico*.

Deep learning enabled ground-breaking performance on tasks across a broad domain of research, including the computational modeling of protein-RNA interaction [1, 14, 5, 41]. Current state-of-the-art RBP binding predictions models are generally classification-based, that is, given an input RNA sequence, the model is tasked with predicting whether the sequence is *bound* or *unbound* by the protein of interest. While classification-based models represent a significant improvement over traditional methods, they have several limitations. First, training and evaluation of classification-based models requires prior annotation of sequences with high-quality binary bound/unbound labels, generally through the use of peaks callers, making the model heavily dependent on upstream preprocessing steps. Therefore, performance of classification-based models is highly sensitive towards the availability of unbiased bona fide binding sites. Second, predictions of classification-based models generally have low resolution. While methods commonly take as input RNA sequences of 100s of nucleotides in length, the predicted label is assigned to the entire input region. This create ambiguity with regards to the exact location of the protein-RNA interaction site, which usually spans only a few nucleotides. Third, binary labels (*bound* and *unbound*) modeled by classification-based methods represent a strong simplification of the information yielded by CLIP-seq experiment. Compression of the CLIP-seq signal footprint within transcript region to a binary value may lead to loss of essential information for understanding the nuances of protein-RNA interaction.

Recently, a new class of models has emerged, which directly predict experimental signal from genomic sequences [2, 27]. For instance, BPNet [2] trains a dilated convolutional neural network which models transcription factor (TF) binding by predicting ChIP-nexus signal from DNA sequences at base resolution. However, there is a lack of sequence-to-signal models for prediction tasks on RNA sequences, including the task of modeling protein-RNA interaction via CLIP-seq signal prediction. In context of CLIP-seq, the presence of technical biases and cell-type specific RNA abundance pose a challenge with respect to the identification of sequence-mediated binding mechanisms at single-nucleotide resolution. Therefore, models need to account for the fact that the observed signal may partially be observed due to technical biases, rather then protein-RNA interaction of target RBP. This effect may further depend on the RNA sequence context, as is the case for nucleotide-specific crosslinking biases or sample contamination with other RBPs which themselves have certain sequence-depended binding preferences.

To fill this gap we developed RBPNet, a sequence-to-signal dilated convolutional neural network, which learns a direct mapping of RNA sequences to crosslink count signal extracted from CLIP-seq experiments. Given a variable-length RNA input sequence, RBPNet predicts the *distribution* of crosslink counts at single-nucleotide resolution along the input sequence, thereby enabling learning the binding profile of an RBP on a transcripts of arbitrary length. Existing sequence-based models of CLIP data are binary classifiers which necessitate important preprocessing steps, thus loosing the high resolution and the quantitative nature of the data In contrast to current state-of-the-art classifiers, RBPNet does not require a peak caller to produce transcript regions as candidate sites for model training. Instead, it uses a lenient cutoff-based approach to train on all regions that show an enrichment of crosslink counts, thereby making maximal use of signal generated by the underlying experiment. By additionally modeling the background signal of a paired control experiment, RBPNet implicitly learns the bias and protein-specific crosslinking components of the total signal, allowing one to disentangle genuine signal from noise. By training on hundreds of thousands of regions per RBP CLIP experiment, RBPNet reaches high accuracy in predicting RBP binding signal shape on held-out test data across eCLIP, iCLIP and miCLIP experiments. Further, it allows for direct inference of predictions across variable-length sequences and whole transcripts, while outperforming state-of-the-art RBP binding classification models with respect to the identification of crosslink sites derived from PureCLIP [31], a single-nucleotide peak caller. By performing model interpretation with Integrated Gradients, we demonstrate the capability of RBPNet to accurately identify the sequence patterns driving RBP-RNA interactions and enable binding motif discovery. Lastly, we show the high potential of RBPNet in scoring the impact of single-nucleotide genetic variants on RBP binding, and thereby enable prioritization of functional variants.

## 2 Results

### RBPNet predicts crosslink count distribution from RNA sequence

RBPNet is a deep convolutional sequence-to-signal neural network for modeling protein-RNA interaction profiles, which takes as input a RNA sequence and outputs a probability vector of the same length, describing a discrete distribution of counts within that sequence. In the context of eCLIP, iCLIP or other individual nucleotide CLIP technologies, RBPNet predicts the distribution of cDNA truncation events, as a result of protein-RNA crosslinking and hereafter also referred to as “crosslink counts”, along an input RNA sequence. In this study, RBPNet was trained and evaluated on a large cohort of eCLIP, iCLIP and miCLIP datasets. An outline of the study is shown in Figure 1a, which gives a schematic overview of training data generation, model training and interrogation for investigating sequence determinants of RBP binding, including binding motif extraction.

**Figure 1:**
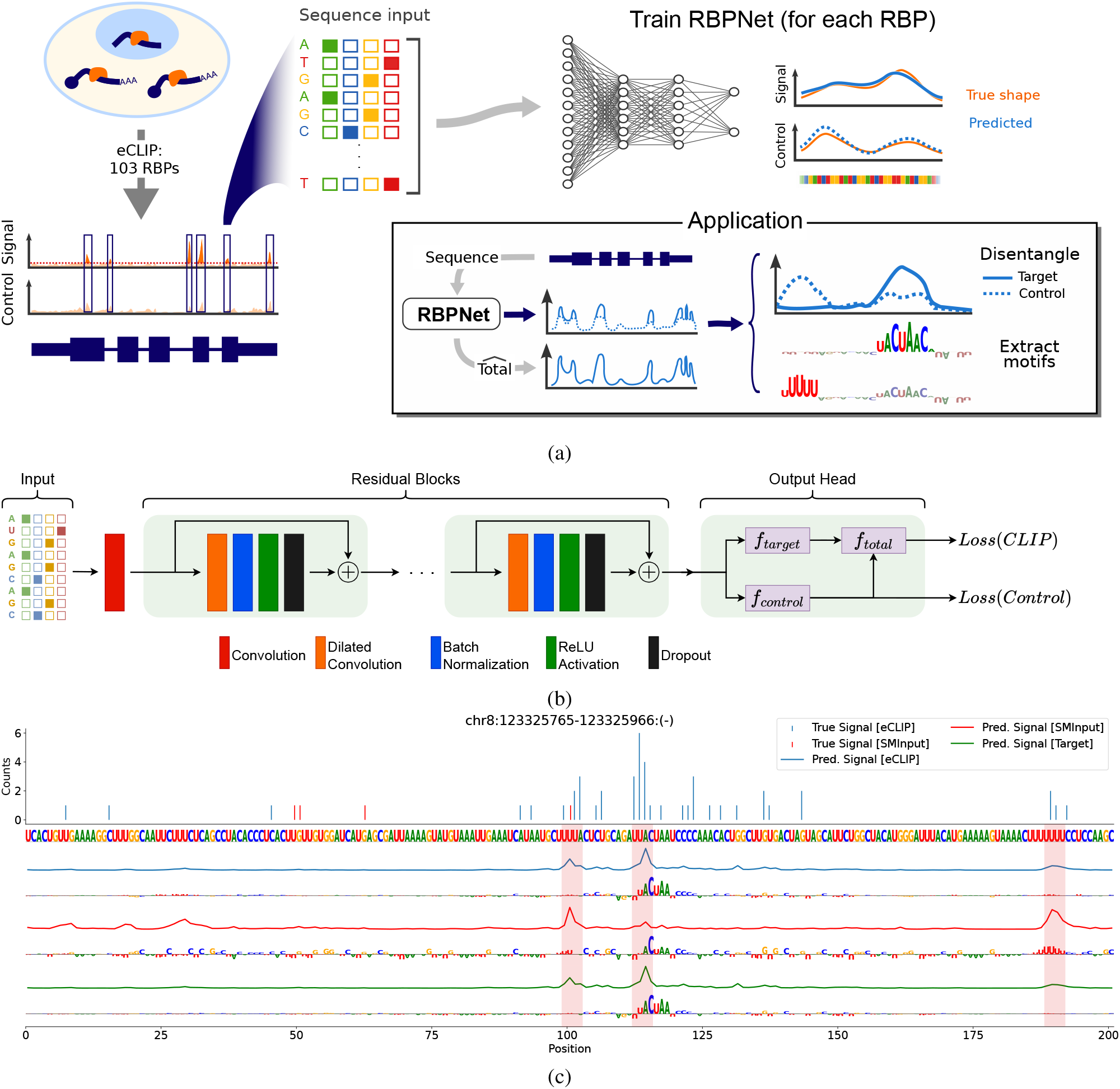
RBPNet overview. **A** Schematic outline of data preparation, RBPNet training and downstream applications. **B** RBPNet model architecture. The one-hot encoded RNA input sequence is first passed through a 1D convolutional layer, followed by several residual blocks, each consisting of a dilated convolution, batch normalization, ReLU and dropout, respectively. Probability vectors of the *target* and *control* tracks are predicted from the output of the last residual block via a transposed convolutional layer while the *total* track is given by an additive mixture of *target* and *control* tracks. Given the predictions, a loss is computed by taking the sum of the negative log-likelihoods of the observed total and control counts. **C** Example prediction of an RBPNet model trained on eCLIP data of QKI showing observed counts (top) and predicted count distributions for the total (blue), control (red) and target (green) tracks. Integrated gradients feature attribution maps with respect to each predict track are shown below.

The RBPNet model architecture consists of two major parts - the model body, comprised of the input layer followed by several convolutional blocks with residual connections, and the model head, which performs the final mapping of the input sequence representation, derived from the body model output, to a probability vector (Figure 1b). Importantly, while RBPNet is trained on fixed-length inputs, its purely convolutional architecture enables prediction on RNA sequences of arbitrary length. During training, the predicted probability vector is used to parameterize a multinomial distribution of crosslink counts and, given the position-wise observed counts in the input sequence interval, a negative log-likelihood loss is computed. In other words, the model is penalized in cases where it is unlikely that the observed crosslink counts were drawn from the distribution predicted by the model. RBPNet thus learns the shape of the crosslink count signal, which is subject to the RNA sequence under the assumption that RNA sequence composition is a driver of recognition (and subsequent binding) by RBPs. Similar to other CLIP-based protocols, eCLIP is known to be subject to experimental biases, for instance as a result of enhanced photoreactivity of single-stranded uridine (U) nucleotides during UV-radiation or contamination of eCLIP libraries with other RBPs [15]. Importantly, these biases are sequence-dependent and directly affect the distribution of cDNA truncation counts, hindering the identification of genuine sequence determinants of RBP binding. For that reason, eCLIP experiments are paired with a size-match input (SMInput) control experiment which omits the protein-specific immunoprecipitation (IP) step, therefore capturing background crosslinking signal from other RBPs or technical biases. To prevent pattern learning of unspecific background and bias signal, RBPNet models the crosslink signal of the control experiment alongside the eCLIP signal. Specifically, RBPNet attempts to explain the observed eCLIP signal as a mixture of two signal components - the *control* component, which is explicitly learned from the control experiment, and an unobserved *target* component, which represents the protein-specific signal (Methods 5.6). The *total* signal, that is, the count distribution of the eCLIP experiment, is then given by a weighted sum of the two components. This is illustrated schematically in the RBPNet output head in Figure 1b as well as in Supplementary Figure 1a, which outlines the network forward pass in the output head. The mixture of the two signals (target and control) is parameterized by a coefficient *π*, which is predicted from the RNA sequence and ranges between 0 and 1. Importantly, we control for technical bias in CLIP assays which we modeled with an additive mixture. This is in contrast to BPNet, a model for chromatin-immunoprecipitation assays which is dominated by DNA accessibility and sequence preference biases that were modeled as multiplicative noise [2].

### RBPNet disentangles bias and protein-specific signal

The formulation of the *total* eCLIP signal as an additive mixture allows for disentanglement of the *target* from the *control* signal, where the predicted *target* signal represents in theory the bias-free, protein-specific crosslinking signal. This is exemplified in Figure 1c, which shows the observed eCLIP and SMInput read start counts (top), as well as the *total, target* and *control* signal predictions (bottom) using an RBPNet model trained on eCLIP data from the QKI RBP. The RBPNet *total track* (blue) captures well the experimental eCLIP read-start count profile, where the highest enrichment of eCLIP counts can be observed at position 113 of the sequence, immediately upstream of the known QKI binding motif (U)ACUUA [6]. Disentangling the predicted signal with RBPNet shows that the enrichment at this position is mostly attributed to the target, i.e. the true QKI-specific binding signal (green). On the other hand, the experimental eCLIP profile harbors two regions with lower enrichment of read start count around relative positions 102 and 189. Disentangling of the RBPNet total signal reveals that the count enrichment in these regions likely originated from experimental bias, as these regions coincide with elevated signal predictions of the control track (red). Further investigation of RBPNet predictions via Integrated Gradients (IG) [50] feature importance scores with respect to each signal track revealed that the known QKI binding motif (U)ACUUA [6] is correctly recovered in the IG map of the target track, corroborating the evidence that the predicted target signal shape corresponds to the bona fide QKI binding signal. In contrast, no clear^2^ QKI motif is observed in the IG maps of the control track, while the recovery of U-rich sequence motifs at the modes of the predicted control track distribution further strengthen the observation that those regions correspond to experimental bias. While the predicted *total* track is a weighted average of *target* and *control* signal (with a mixing coefficient of 0.92, such that the target is dominating over the control track), genuine QKI binding signal is correctly recovered by the predicted *target* track. Likewise, signal enrichment mainly representing experimental bias are recovered by the *control* track, while being present in very low proportions in the *target* signal.

### RBPNet predicts eCLIP signal shape at replicate-level accuracy

We next performed evaluation on 103 RBPNet models trained on data from ENCODE [53] eCLIP experiments. To this end, for each eCLIP experiment, candidate training pairs of 300nt long sequences and crosslink count footprints were first generated without the use of a peak caller via lenient signal-thresholding (Methods) and subsequently split chromosome-wise into train, validation and test sets (Methods). Overall, we obtained an average of 302, 752 candidate sites across eCLIP datasets, with a minimum of 7, 937 sites for LARP7 and a maximum number of 1, 105, 807 sites for HNRNPC. Subsequently, RBPNet models were trained separately for each RBP for at most 50 epochs, while the validation loss was observed for the purpose of early-stopping (Methods). Example predictions on hold-out samples with highest total observed counts for TIA1, QKI and U2AF2 are depicted in Figure 2a. Predictions show a high Pearson correlation coefficient (PCC) with the observed eCLIP counts (0.763, 0.830 and 0.872 for TIA1, QKI and U2AF2, respectively), demonstrating that RBPNet can recapitulate the eCLIP signal shape at high accuracy. Next, we quantitatively assessed correlation of predicted and observed signal distribution. Figure 2b shows the average PCC of RBPNet (total track) predictions with observed eCLIP counts on hold-out samples versus the average PCC of counts between the two eCLIP replicates on the same sequence intervals. RBPNet achieves an average PCC between 0.200 (SSB) and 0.587 (HNRNPC), with an average PCC of 0.328. Strikingly, RBPNet prediction appear to outperform replicates, which have an average PCC of 0.149 across all RBPs. The reason for this are two-fold. First, correlation between RBPNet predictions and observed counts are computed with respect to the merged count of both replicates, which reduces sampling effects and thus may increase PCC. Second, RBPNet predicts the count-generating distribution in the given interval, conditioned on the RNA sequence. In contrast, the observations in each replicate represent samples from the true (but unknown) count-generating distribution. As the estimated signal distribution by RBPNet approaches true distribution, the expected PCC between RBPNet predictions and a sample exceeds the expected PCC between the two samples (Supplementary Text).

**Figure 2:**
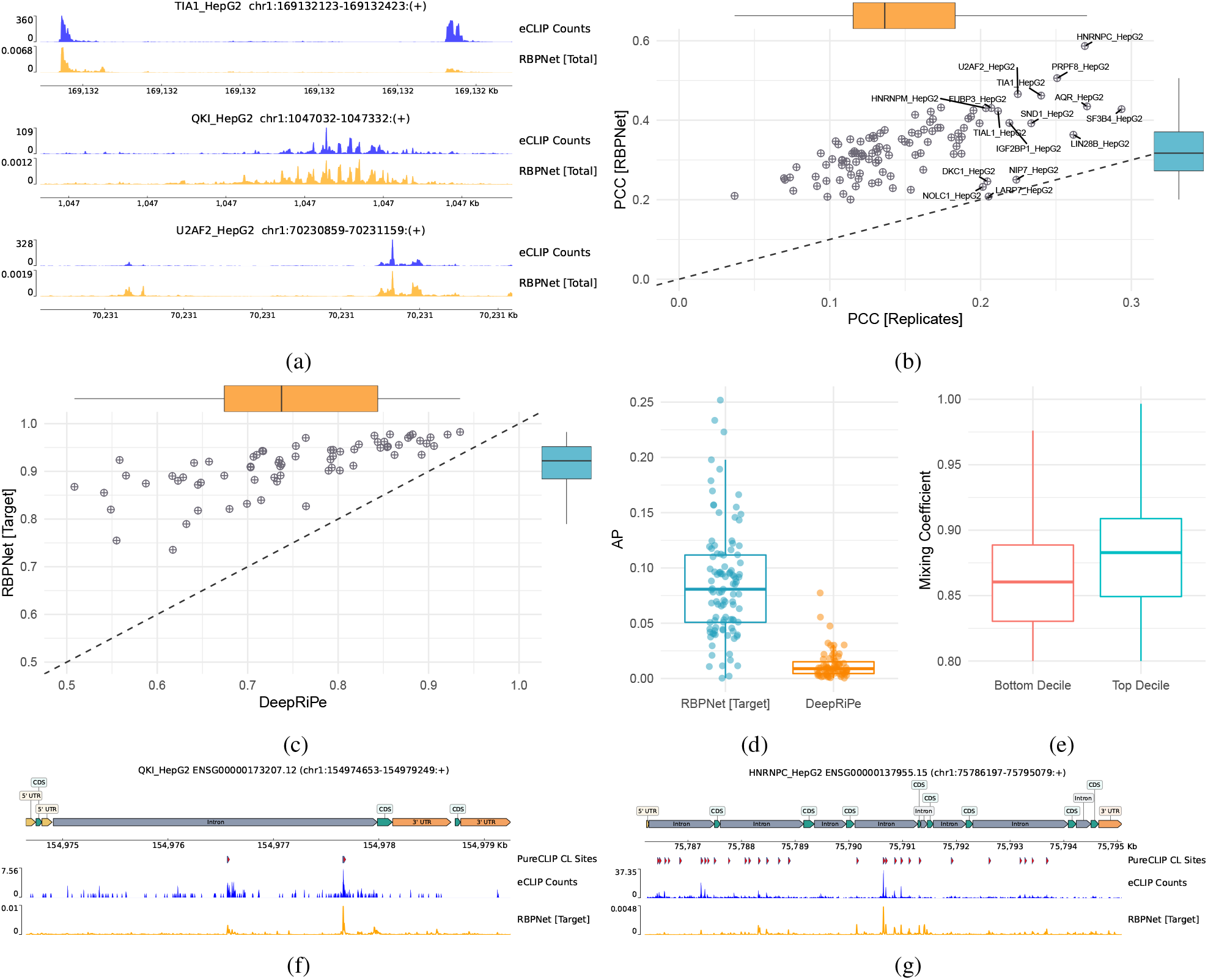
RBPNet prediction performance on ENCODE eCLIP datasets. **A** RBPNet predictions (total track) on the highest-count hold-out samples for TIA1, QKI and U2AF2. **B** Pearson correlation coefficient (PCC) of RBPNet predictions (total track) with observed eCLIP crosslink counts on hold-out samples vs. PCC of observed counts between the two eCLIP replicates. **C, D** Mean auROC and AP of RBPNet (target track) and DeepRiPe predictions with respect to crosslink and non-crosslink positions called by PureCLIP across transcripts from hold-out chromosomes, respectively. As pre-trained DeepRiPe models are available only for 70 (out of 103) ENCODE HepG2 RBPs, performance comparison is shown only for those RBPs. **E** Distribution of RBPNet mixing coefficients of the top and bottom decile ENCODE narrow peaks, sorted by eCLIP signal fold-change over the SMInput. High-affinity ENCODE narrow peaks show on average higher mixing coefficients compared to low-affinity peaks. **F, G** Example RBPNet (target track) whole-transcript predictions on ENSG00000173207.12 and ENSG00000137955.15, together with observed eCLIP counts and called PureCLIP peaks, for QKI and HNRNPC, respectively.

### RBPNet enables whole-transcript inference and recovers single-nucleotide resolution binding sites

We next leveraged RBPNet’s ability of performing prediction of RNA sequences of arbitrary length, despite being trained on fixed-length inputs. We explored first whether RBPNet can infer signal on entire transcripts by first selecting genes from GENECODE (Release 40) [10] from hold-out chromosomes and subsequently performing RBPNet predictions using models for all ENCODE RBPs. Figure 2f and 2g shows RBPNet predictions (total track) on ENSG00000173207.12 and ENSG00000137955.15 for QKI and HNRNPC, respectively. Indeed, RBPNet predictions show a high correlation with the observed eCLIP counts (0.645 and 0.816, respectively), demonstrating that RBPNet models trained on rather short, fixed-size inputs generalize well to the task of whole-transcript prediction.

Given that RBPNet predictions show high correlation with observed eCLIP counts, we next assessed whether high-scoring RBPNet predictions coincide with peaks called by PureCLIP [31], a single-nucleotide peak caller that identifies significant crosslink sites from eCLIP and SMInput cDNA truncation counts using a Hidden Markov Model. To this end, we performed whole-genome peak calling with PureCLIP on ENCODE eCLIP datasets (Methods 5.10.2), identifying on average 46, 459 crosslink (CL) sites per RBP, with a minimum and maximum number of CL sites of 1, 083 and 585, 772 for NIP7 and HNRNPC, respectively. For each RBP, hold-out chromosome genes were intersected with PureCLIP CL sites and transcripts harboring at least 10 CL sites were selected for downstream evaluation. Subsequently, whole-transcript signal shape prediction was performed via RBPNet on selected transcripts. Here, predictions of the *target* track were used as PureCLIP peaks were called using both eCLIP and SMInput background signal information (Methods). For each transcript, auROC and average precision (AP) performance metrics were computed by treating positions at PureCLIP CL sites as positives and all other positions as negatives (Supplementary Table 1). Since RBPNet predicts the distribution of crosslink sites along genes, the probability at each position correlates with the genes length as well as the abundance of RBP binding on the transcript, rendering position-wise RBPNet predictions uncomparable across transcripts. Therefore, auROC and AP metrics are computed within transcripts and later averaged across transcripts in order to report the final, RBP-specific auROC and AP scores. In order to assess how the RBPNet behaves with respect to classification-based models, we next compared RBPNet predictions to DeepRiPe, a state-of-the-art classifier for prediction of protein-RNA interaction, by using a sliding-window approach to obtain pseudo single-nucleotide resolution scores (Methods). Since pre-trained DeepRiPe models for HepG2 ENCODE datasets were obtained directly from Ghanbari et al. [14], sequences in our hold-out set may have been present during training of DeepRiPe.

Figure 2c and 2d show the average auROC and AP scores, respectively. RBPNet outperforms DeepRiPe on all RBPs in terms of auROC performance, with an average auROC of 0.89 and a minimum and maximum auROC of 0.58 and 0.98 for SSB and HNRNPK, respectively. In contrast, DeepRiPe achieves a significantly lower average auROC of 0.74. Interestingly, RBP-wise auROC scores of RBPNet and DeepRiPe are strongly correlated (PCC = 0.72). This may suggest that some ENCODE eCLIP libraries are of lower quality or that RNA binding of some RBPs is more difficult to predict from sequence, possibly due to a lack of sequence binding motifs. Notably, RBPNet shows a lower variance of auROC performance across RBPs. Given that classification-based models such as DeepRiPe rely heavily on proper categorization of RNA sequences into binding and non-binding for training, this may indicate a higher robustness of RBPNet due to its training approach, which does not rely on upstream peak calling or labeling of RNA sequences. Both RBPNet and DeepRiPe AP values across RBPs range between 0.00032 and 0.2518 and .00038 and 0.0774, respectively, with RBPNet significantly outperforming DeepRiPe (average AP of 0.086 vs. 0.012, respectively). The generally low AP values of both methods is due to AP being sensitive to class imbalance, with the random AP baseline being equivalent to the fraction of positive samples in the dataset. Here, hold-out transcripts have an average PureCLIP CL site fraction of 0.0014 across RBPs, given that transcripts are expected to harbor orders of magnitude more non-CL than CL sites. Thus, while having low AP, RBPNet and DeepRiPe outperform the random baseline by a large margin. Overall, these results show that RBPNet is a powerful discriminator of PureCLIP CL and non-CL positions across the transcriptome, outperforming state-of-the-art classifiers.

### RBPNet mixing coefficient captures relative eCLIP and SMInput signal abundance

RBPNet models the *total* eCLIP signal as a mixture of protein-specific *target* and *control* signal, weighted by a mixing coefficient which determines the fraction of *target* signal in the *total* signal (Methods). In order to evaluate whether the mixing coefficient captures the fraction of target and control signals properly, we inspected mixing coefficients of high-and low-affinity ENCODE narrow peaks [54]. To this end, for each RBP, we obtained ENCODE narrow peaks and ordered them decreasingly with respect to their log2 fold-change (logFC) of eCLIP signal over the SMInput. Peaks in the top and bottom deciles were then selected and extended up-and down-stream from their 5’ end, which generally corresponds to the crosslink site, to yield 300nt windows. Finally, RBPNet predictions were performed for each window and mixing coefficients were obtained. Figure 2e shows the distribution of mixing coefficients on top and bottom decile ENCODE narrow peaks. Indeed, top decile peaks receive on average significantly higher mixing coefficients compared to bottom decile peaks (*p <* 2.2 × 10^−16^), suggesting that the RBPNet mixing coefficient can separate high-affinity from low-affinity sites, where the latter contains a higher proportion of background signal.

### RBPNet generalizes to iCLIP and miCLIP experiments

RBPNet may be trained on any genomic sequence with a corresponding nucleotide-wise count signal. To demonstrate that RBPNet generalizes to other CLIP-based protocols, we trained RBPNet on data derived from miCLIP and iCLIP experiments.

miCLIP enables *in vivo* identification of m6a RNA modifications at single-nucleotide resolution by incubating and subsequently crosslinking extracted RNA with a m6a-specific antibody [36]. After digestion of the covalently bound antibody, reverse transcription often truncates at a remaining polypeptide at the crosslink site, with pileups of truncation events yielding an m6a count signal across the transcriptome. miCLIP data from HEK293 and mESC cells was gathered from Kortel et al. [30] and processed similar to eCLIP data (Methods), before selecting candidate sites for training and evaluation (Methods). Subsequently, RBPNet models were trained on both cell lines using a similar architecture and hyperparameters as in eCLIP RBPNet models (Methods). While the mESC miCLIP experiment was paired with a knockout (KO) control experiment, no control experiment was available for HEK293. We therefore trained RBPNet on HEK293 miCLIP data without *target* and *control* modeling, that is, RBPNet was tasked to predict a single track describing the distribution of *total* miCLIP counts in the HEK293 cell line. RBPNet (total track) showed high signal correlation on miCLIP counts in both cell lines (Figure 3a), reaching PCCs of 0.51 and 0.48 for HEK293 and mESC, respectively. To evaluate the ability of RBPNet to recover miCLIP single-nucleotide peaks, we again performed peak calling with PureCLIP (Methods), yielding a total of 2, 011, 704 and 278, 311 for HEK293 and mESC miCLIP, respectively. We hypothesize that the significantly higher number of PureCLIP CL sites in HEK293 may be due to a lack of control signal, which in context of mESC may lead to a large number of candidate CL sites being discarded. While the KO control is used for PureCLIP background normalization for the mESC cell line, no control is used for peak calling on the HEK293 cell line, as this was not available. Due to the lack of controls, PureCLIP yields a significantly higher number of CL sites in HEK293 compared to mESC, where a high portion of them may correspond to false positives or noise. Similar to PureCLIP analysis in eCLIP, we then performed whole-transcript inference of miCLIP signal shape on genes of the hold-out set. Figure 3b and 3c show the ROC curves for HEK293 and mESC, respectively. RBPNet performs well in terms of auROC on both cell lines, with an average auROC of 0.89 for HEK293 and 0.88 for the mESC cell line. In contrast, RBPNet achieves an AP scores of 0.1 and 0.043 for HEK293 and mESC, respectively, with AP performance naturally increasing together with the fraction of positions harboring PureCLIP CL sites. This is illustrated in Figure 3d, which shows the distribution of AP scores across transcripts grouped into quartiles based on their CL fraction. We consequently observe a lower AP score for mESC compared to the HEK293 cell line as a result of the lower number of PureCLIP CL sites in mESC. Figure 3e and 3e shows example RBPNet predictions (total track) on two human genes (ENSG00000142937.11 and ENSG00000161016.15, respectively) from the hold-out chromosomes using the HEK293 RBPNet model. Interestingly, RBPNet predictions (as well as observed signal) predominantly occur in coding regions (CDS) and 5’/3’ untranslated regions (UTRs), while being only slightly present in introns, which is in line with previous study reporting that *m*^6^*A* methylation mainly those genomic regions [26, 38].

**Figure 3:**
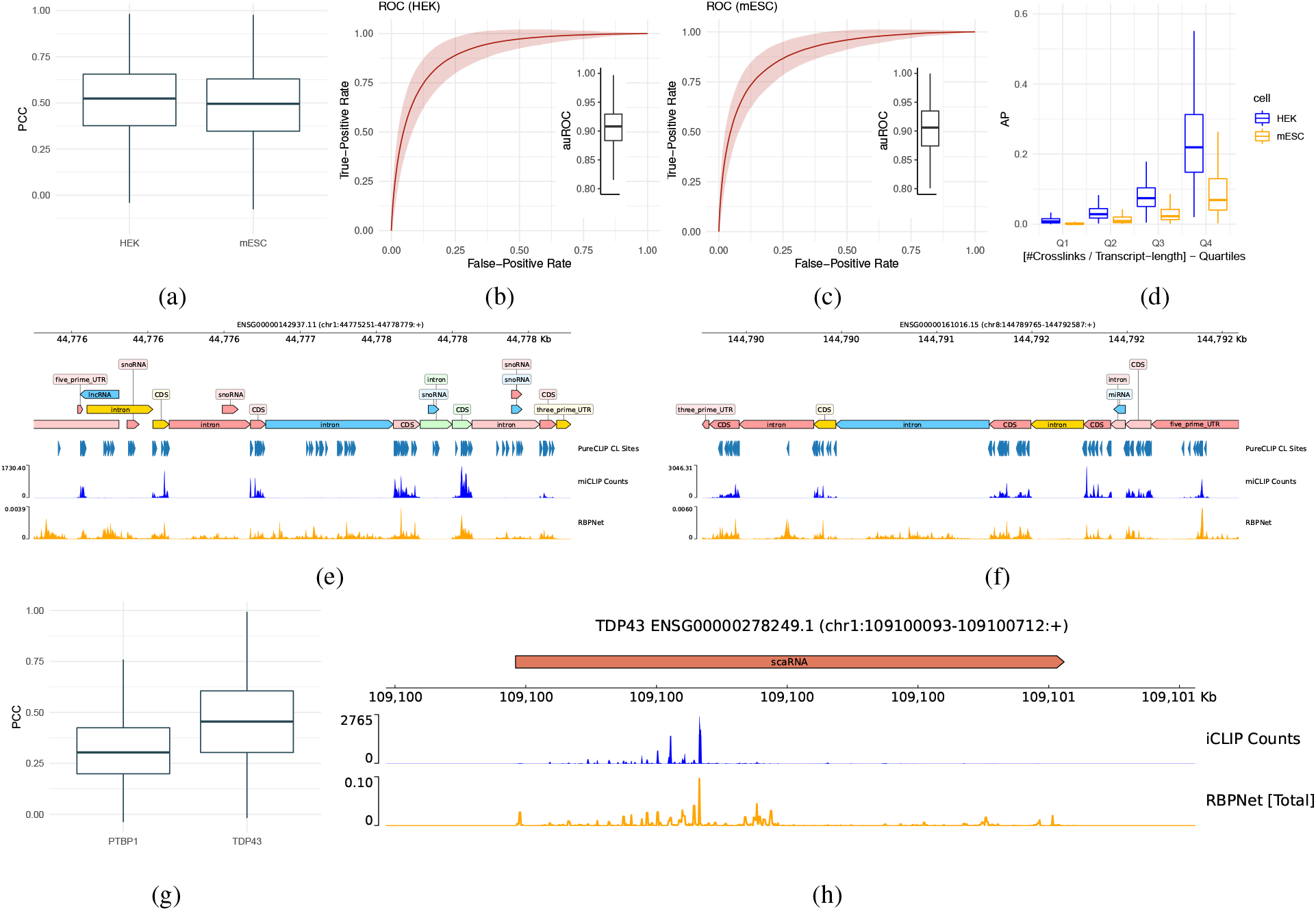
Evaluation of RBPNet on iCLIP and miCLIP data. As in contrast to mESC, no control signal is used for m6a peak calling on HEK data, we consequently used RBPNet target and total track predictions for auROC and AP computation for mESC and HEK, respectively. **A** RBPNet PCC performance on hold-out samples of miCLIP experiments in HEK293 and mESC cell lines. **B, C** ROC performance of RBPNet whole-transcript predictions with respect to crosslink and non-crosslink positions called by PureCLIP for HEK293 and mESC cell lines, respectively. **D** AP performance on HEK293 and mESC in across transcripts for different PureCLIP crosslink site frequency quartiles. **E, F** Example RBPNet-HEK293 miCLIP predictions on ENSG00000142937.11 and ENSG00000161016.15. Notably, both predicted signal shape and observed miCLIP signal occurs predominantly in CDS and UTR regions. **G** Test set PCC performance on iCLIP experiments for PTBP1 and TDP43. **H** Predicted RBPNet signal shape for SCARNA2 (ENSG00000278249.1).

We next evaluated the performance of RBPNet on individual-nucleotide CLIP (iCLIP) data [29]. Compared to eCLIP, it makes use of extra circularization and linearization steps, which allow all cross-linked cDNA fragments to be amplified and sequenced, and a quality control step to assess specificity of pulled-down protein-RNA complexes. To this end, we gathered iCLIP data from Hallegger et al. [18] and Haberman et al. [15] for TDP43 and PTBP1 proteins, respectively, which was processed as described in Section 5.1. As no paired control experiment was available for either RBP, we again trained RBPNet by omitting modeling of *target* and *control* signal and instead tasked RBPNet with predicting the total iCLIP count distribution directly. Figure 3g shows the distribution of PCCs on test set samples for the PTBP1 and TDP43 models. RBPNet reaches an average PCC of 0.46 for TDP43 and 0.320 for PTBP1. Notably, RBPNet reaches a comparable PCC of 0.366 on the PTBP1 eCLIP dataset, which may suggest that the fraction of PTBP1 signal (and thus RNA-binding) explained by RNA sequence is comparable in the eCLIP and iCLIP datasets. This demonstrates the capability of RBPNet to achieve high predictive performance of the RBP-RNA interaction signal shape, independently of the protocol used to generate the data the model is trained on. In order to qualitatively assess the ability of the RBPNet-TDP-43 model to predict signal shapes on full length transcript when trained on iCLIP data, we manually investigated the prediction profile on the hold-out transcript with the highest absolute counts. Figure 3h shows TDP-43 RBPNet predictions and observed iCLIP counts for ENSG00000278249.1 (SCARNA2), a scaRNA associated with DNA repair pathway regulation that has been previously described to be interacting with TDP43 [23, 4]. Indeed, the profile predicted by RBPNet strikingly reflects the observed signal.

### Sequence attribution maps capture RBP binding motifs

Recognition of target RNAs by RBPs is in part driven by local sequence features, also known as binding motifs. The identification of RBP binding motifs is crucial for understanding RBP target recognition and the regulatory grammar present in RNA sequences. While deep learning models were long regarded as black boxes, recent feature attribution methods, such as Integrated Gradients (IG) [50], allow for the identification of input features that contributed significantly to the observed model prediction. In the context of RBPNet, these methods “attribute” a given prediction to nucleotides in the input RNA sequence by assigning a score to each position. Nucleotides that were primarily responsible for the observed crosslink count distribution, such as those residing in binding motifs, receive a higher score compared to nucleotides that did not contribute towards protein-RNA crosslinking. IG attribution maps may be computed with respect to any of the three output track, i.e. control, target and total. Attributions of the target track are expected to highlight nucleotides that contributed significantly towards protein-specific crosslinking, while attribution of the control track are expected to explain the unspecific background and bias signals^3^.

Figure 4a shows examples of IG attribution maps, computed with respect to *control* and *target* tracks for test-set samples using eCLIP RBPNet models for RBFOX2, HNRNPK, TIA1 and QKI, alongside PWMs of consensus motifs reported in literature, obtained from the RBPmap database [43]. The corresponding predicted *control* and *target* signals are shown above the attribution maps (in red and green, respectively). Target track IG attribution maps show the presence of highly predictive sub-sequences that correspond to known binding motifs for each of the shown RBPs. For instance, IGs of HNRNPC show the characteristic U-motif, while three distinct canonical ACUAAC motifs can be seen in the IGs of QKI. In contrast, control track attribution maps do not show the presence of clear binding motifs, with a general attribution score enrichment at G and C nucleotides. We next performed a global quantitative assessment of how well RBPNet attribution maps can recover known binding motifs across RBPs in the ENCODE eCLIP database. To this end, we selected the top 5, 000 ENCODE narrow peaks for all ENCODE eCLIP experiments with trained RBPNet models and computed IG attribution maps with respect to *target, control* and *total* tracks for each RBP. For each attribution map, we extracted 5-mers with highest sum IG scores (Methods). Extracted 5mers were then compared with position-weight matrices (PWMs) of literature motifs obtained from the RBPmap [43] database by computing the similarity between each 5mer and its corresponding RBPmap motif (Methods). In total, 29 out of the 103 ENCODE RBPs with RBPNet models were represented with at least one PWM in RBPmap, which were then selected for downstream analysis. Figure 4b shows the distribution of average similarity scores of extracted 5mers to RBPmap PWMs across RBPs for each of the three RBPNet prediction tracks. 5-mers extracted from attribution maps of the *target* track show the highest RBPmap-similarity, consistent with the fact that this track represents the de-biased, protein-specific signal predictions, followed by the *total* and *control* tracks, respectively. Notably, on average 5-mers extracted with respect to the control track have the lowest similarity to known RBP PWMs, highlighting the ability of RBPNet to extract different sequence representation for true signal and bias, respectively. To investigate the degree of improvement that signal de-biasing in the *target* track offers over the *total* track for individual RBPs, Figure 4c shows the average *target* and *total* track RBPmap-similarity for each RBP. Interestingly, we find that while for some RBPs, the *target* track leads to no or only modest improvements of RBPmap-similarity compared to the *total* track, other RBPs, such as HNRNPC, PCBP2, RBFOX2 and TRA2A, appear to benefit strongly from the signal de-biasing of the *target* track. This may reflect variable levels of experimental bias across eCLIP datasets.

**Figure 4:**
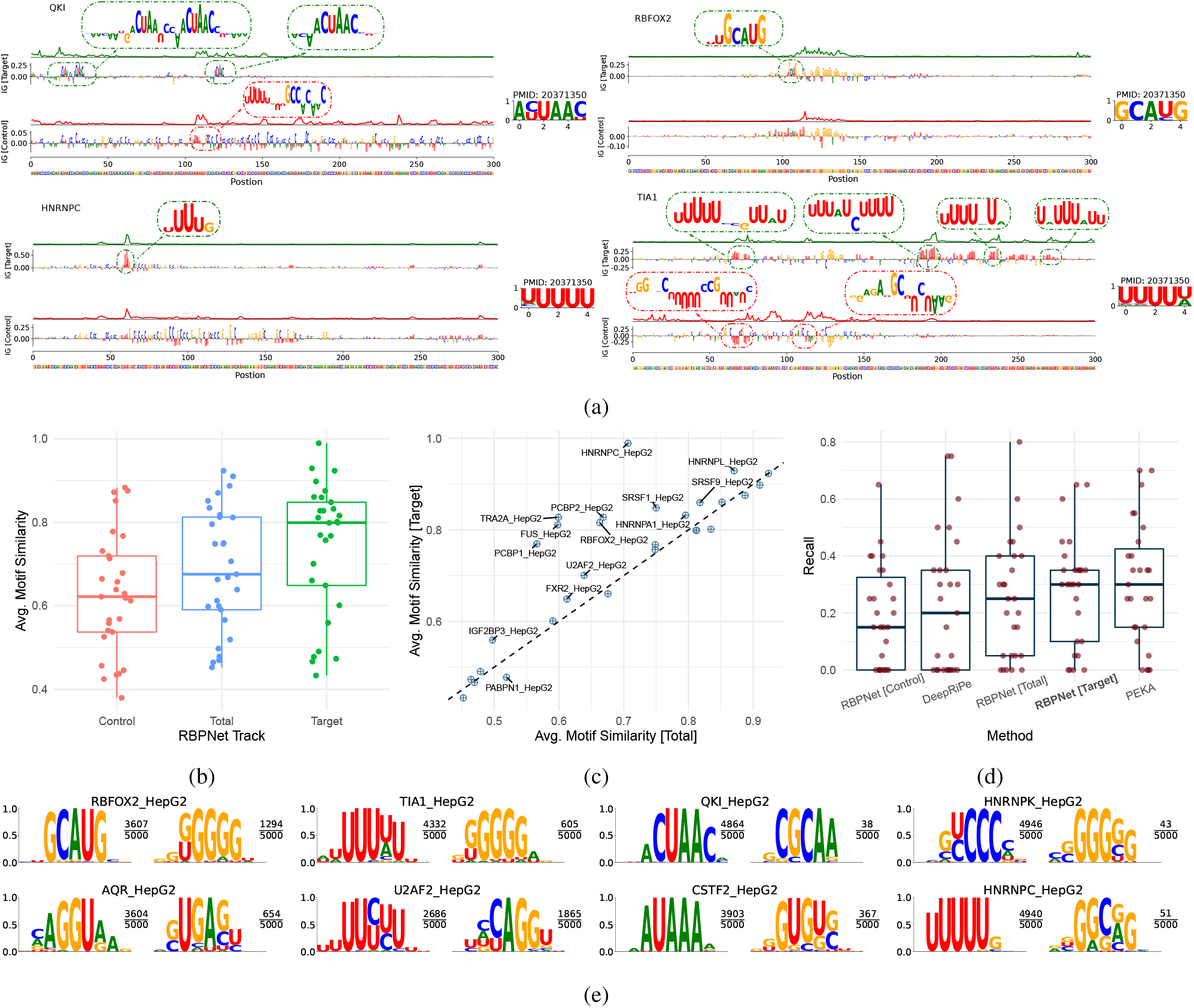
RBPNet feature attribution maps and binding motif discovery. **A** Example integrated gradient attribution maps with respect to the *target* track for RBFOX2, HNRNPK, TIA1 and QKI with corresponding motifs taken from the RBPmap database. **B** Distributions of similarity scores between 5-mers extracted from RBPNet attribution maps and PWMs of motifs reported in RBPmap for *control, target* and *total*. **C** Average RBPmap PWM similarity across RBPs for 5-mers extracted from *target* and *total* track attributions. While similarities of the *target* track on par or higher compared to the *total* track for the vast majority of RBPs, the improvement is more pronounced for some RBPs. **D** Recall of the top 20 *in vitro* 5-mers recovered by the top 20 5-mers extracted from attribution maps of RBPNet *control, target* and *total* tracks as well as DeepRiPe and the PEKA motif finder. RBPNet *target* track and PEKA show comparable performance, outperforming DeepRiPe and the RBPNet *total* track. **E** Consensus motifs constructed from extracted 5-mers of the RBPNet *target* track. Consensus motifs with highest (primary) and second highest (secondary) k-mer support are shown. The corresponding k-mer support is shown as a fraction of the total number of extracted 5-mers next to the consensus motif logos.

### RBPNet IG attribution maps recover *in vitro* binding motifs

*In vitro* experiments on protein-RNA interactions, such as RNA-Bind-N-Seq and RNAcompete, offer an orthogonal view to validate motifs identified from *in vivo* data, as they do not harbor crosslinking specific biases or contamination of experiments with other RBPs and therefore measure intrinsic affinity of RBPs to RNA in an unperturbed environment [34, 45]. To examine whether modelling of the control signal as an auxiliary task can increase the specificity of predicted CLIP signal, we cross compared 5-mers previously obtained from RBPNet IG attribution maps of *target, control* and *total* tracks with 5-mers enriched in corresponding RNA-Bind-N-Seq (RBNS) or RNAcompete *in vitro* datasets. In total, we evaluated 27 eCLIP datasets in the HepG2 cell line, for which either RBNS (16 RBPs) or RNAcompete (11 RBPs) data was also available.

To measure the agreement between *in vitro* 5-mers and RBPNet 5-mers obtained in the previous section, we first computed the sum of *IG*_*sum*_ scores for each unique 5-mer as a measure of importance with respect to CLIP signal shape prediction. We then calculated the RBPNet recall for each track by taking the fraction of the top 20 *in vitro* 5-mers that were recovered in the top 20 5-mers by RBPNet on eCLIP datasets (Methods). We found that across evaluated RBPs, the RBPNet *target* track recovered a significantly higher proportion of relevant *in vitro* 5-mers from eCLIP than the *total* track, suggesting that RBPNet can successfully increase the specificity of eCLIP signal (Figure 4d). As expected, the *control* track recovered the least *in vitro* k-mers, however, for some RBPs even the *control* track alone could retrieve high ranking *in vitro* motifs. This effect could be explained by a partial enrichment of RBP-specific signal in the control experiment, as suggested by a previous study, which evaluated the effect of using eCLIP narrow peaks in contrast to SMInput-agnostic peak-calling on discovery of relevant binding motifs from eCLIP data [32]. Next, we set out to assess whether the governing sequence features learned by RBPNet could be reliably used for motif discovery. To this end, we compared the RBPNet recall to positionally-enriched k-mer analysis (PEKA), a state-of-the art tool for discovery of enriched k-mers from individual CLIP datasets [32] and DeepRiPe. In contrast to RBPNet, PEKA models background from intrinsic crosslinking signal and does not use SMInput controls, therefore it provides valuable orthogonal view in how specificity of the motifs can be impacted by different background modelling approaches. Surprisingly, we found that the RBPNet *target* track recovered a similar proportion of relevant *in vitro* k-mers compared to PEKA, despite the fact that RBPNet was not originally designed for objective of motif discovery (Figure 4d). Lastly, we investigated whether a direct modeling of eCLIP signal, compared to binary labels (bound / unbound) assigned to entire RNA sequences, offers an advantage in the context of motif discovery. To this end, we compared RBPNet recall performance to DeepRiPe by extracting 5-mers from DeepRiPe attribution maps (Methods). Indeed, both RBPNet *target* and *total* track outperform DeepRiPe in terms of *in vitro* k-mer recovery (mean recall of 0.294, 0.254 and 0.222, respectively), suggesting that direct modeling of raw eCLIP signal improves binding motif discovery irrespectively of bias modeling. Further, these results indicate that enriched *in vitro* k-mers are strong predictors of true CLIP signal and that RBPNet learns to associate the presence of these k-mers with high count signal. Lower agreement of the RBPNet *total* track (compared to *target* track) with *in vitro* k-mers suggests that a fraction of the *total* track signal is explained by sequence features not present in *in vitro* k-mers, which likely correspond various possible technical sources of sequence biases or contaminant signals in CLIP [16, 32].

### Consensus motifs reveal primary and secondary motifs

Having demonstrated that RBPNet successfully identifies predictive k-mers that coincide with *in vitro* datasets, we next constructed consensus binding motifs to concisely represent the sequence binding preference of each eCLIP RBP. To this end, we select 5-mers previously derived from RBPNet *target* track attribution maps and build consensus motifs by successively aligning 5-mers in the order of highest to lowest IG importance (Methods). Figure 4e shows consensus motifs together with the fraction of supporting 5mers for selected RBPs. Primary binding motifs identified with RBPNet agree with previously identified motifs reported in RBPmap (RBFOX2, QKI, TIA1, U2AF2, HNRNPC). Additionally, we identified novel candidate motifs for several RBPs, including AQR and CSTF2, for which no motifs exist in the RBPmap database. We further identify secondary consensus motifs, which may represent alternative binding preferences or co-factor binding (Methods). Several RBPs, including RBFOX2, TIA1 and (to a lesser extend) HNRNPK, show a G-rich secondary motif (Figure 4e). We hypothesize that this may be due to co-factor binding of an RBP with G-motif preference or, as suggested by previous studies, may indicate contamination of eCLIP libraries with another RBP 32, 53]. A complete list of consensus motifs derived from RBPNet are shown in Supplementary Figure 2.

### Studying the impact of sequence variants on protein-RNA interaction with RBPNet

RBPs are highly evolutionary conserved and have been associated with an abundance of human diseases, particularly in the context of degenerative disorders [42, 7]. Recently, Gebauer et al. [13] found that over 1,000 RBPs are mutated in context of disease, which amounts to >20% of proteins annotated with disease-associated mutations. Besides altered coding sequences of RBPs, nucleotide polymorphisms in their RNA targets may impact transcript regulation via loss of binding sites. Indeed, Park et al. [42] showed that dysregulation of RNA target sites from RBPs with a diverse set of functions represents is a key driver of psychiatric disorder risk. Therefore, computationally quantifying the impact of variants with respect to protein-RNA interaction at large scale is crucial for the prioritization of causal variants in context of disease. Here, we use RBPNet to score the impact of sequence variants on protein-RNA interaction and demonstrate that resulting impact scores yield candidate variant that may be associated with the disruption of protein-RNA interaction.

Figure 5a exemplifies our variant scoring approach on *rs6981405*, an A-to-C transversion within the *DDHD2* gene that disrupts QKI binding and which has been associated with schizophrenic risk [42]. As shown, *in silico* mutagenesis leads to change in predicted RBPNet target signal around the variant, with a lower signal amplitude in the alternative (ALT) allele compared to the reference (REF), which we quantify as the Kullback–Leibler (KL) divergence between REF and ALT predictions (Methods), yielding a scalar KLD impact score. Feature attribution maps (Figure 5a, bottom) of the REF and ALT predictions reveal that *rs6981405* disrupts the well known QKI binding motif ACUAAC [17]. The strongly negative implications of the A-to-C transversion at the last position of the binding motif is correctly detected by RBPNet, as is evident from the change of high-positive to high-negative attribution at the mutation site. To evaluate whether RBPNet assigns greater impact to SNPs within the QKI motif compared to SNPs outside the motif, we performed a systematic *in silico* perturbation analysis of each nucleotides within a 200nt window around the *rs6981405* SNP. Figure 5b depicts the distribution of a total of 600 variant impact scores, grouped based on whether they reside within or outside the QKI binding motif. Indeed, the majority of non-motif perturbations lead to small changes in predicted signal profile and thus to small impact scores, while mutations falling within the QKI motifs lead to significantly larger impact scores. This demonstrates that RBPNet can successfully quantify variant impact through the change of its predicted signal between REF and ALT alleles.

**Figure 5:**
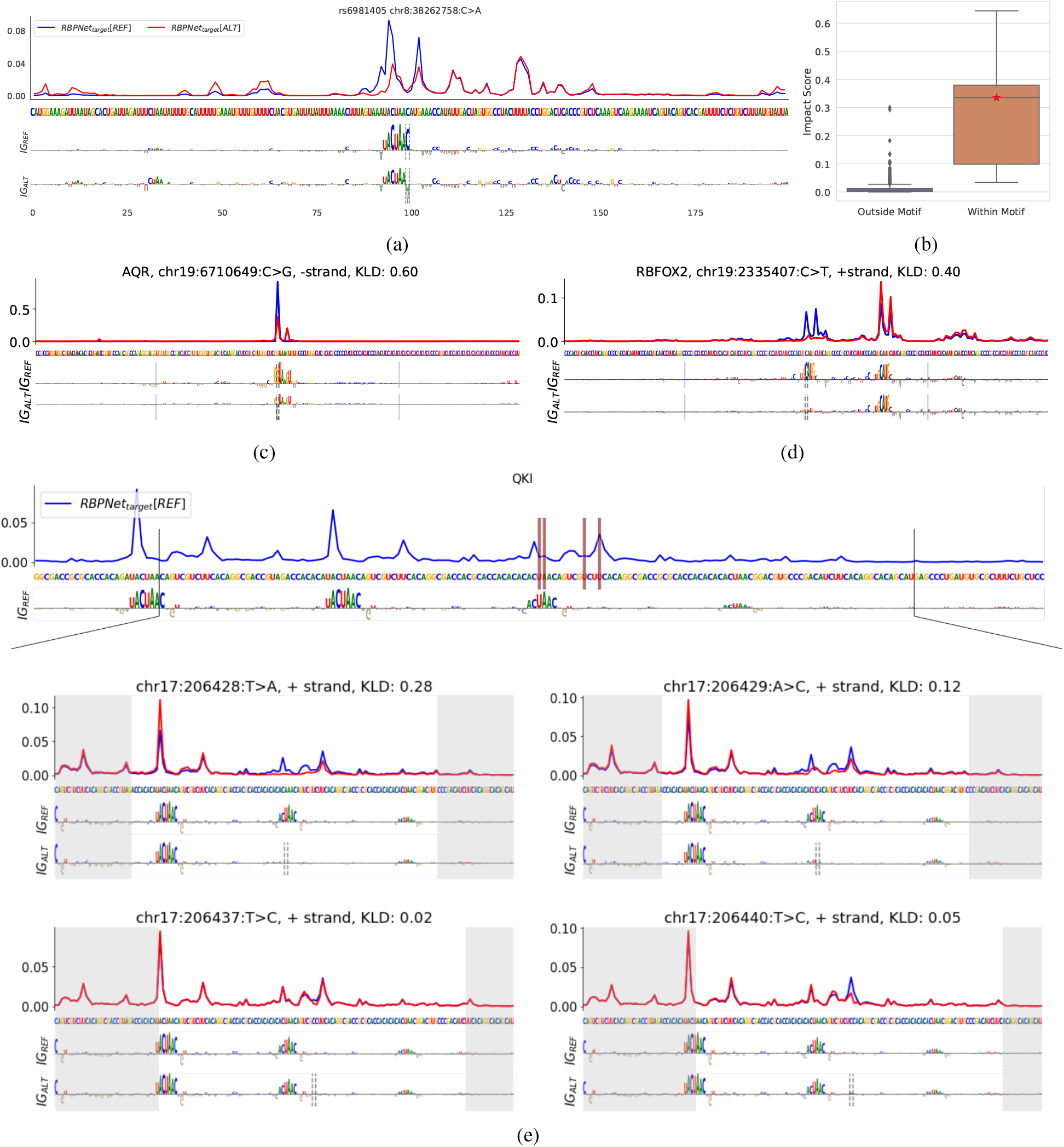
RBPNet variant impact prediction. **A** Impact of the *rs6981405* variant on the predicted RBPNet *target* signal. Reference (blue) and alternative (red) signal is shown in a 200nt window around the variant position. The corresponding feature importance maps for reference and alternative sequence are shown below. A C-to-A transversion at the 5’ end of the QKI binding motif leads to a drastic change of the predicted signal compared to the reference signal. **B** Comparison of impact scores of a systematic *in silico* mutagenesis of each position towards each of the 3 alternative bases within in a 200nt sequence window around *rs6981405*. Note that mutating a nucleotide within the motif leads to a significantly higher variant impact score than outside the motif. The red star indicates the score of the observed rs6981405 variant. For scoring variants we used KL-divergence between the reference and alternative profile. **C, D** Example impact predictions of the most significant allele-specific binding (ASB) events identified by [58] for AQR and RBFOX2. **E** Variant impact predictions for the QKI model in a region harboring 4 distinct SNVs associated with ASB. The top plot shows the reference sequence with the corresponding attribution map. The location of the SNVs within the sequence is highlighted with red bars. The 4 SNVs are displayed separately, showing a comparison of the predicted profile with the reference (top) together with the attribution maps of the reference and the alternative allele (bottom). The location of the variant is highlighted with a dashed grey line in the attribution map of the alternative allele. The grey boxes represent the margins of the KLD scoring window centered around the variant. RBPNet predicts notable allele specific effects for only 2 out of the 4 candidate SNVs, which coincide with QKI binding motifs. These SNVs are associated with a loss of the binding motif in the attribution map and a change in the predicted signal.

### Investigating allele-specific binding (ASB) with RBPNet

To further probe RBPNet’s ability to predict trustworthy variant impacts on protein-RNA interactions, we manually investigated experimentally identified allele specific binding events report by Yang et al. [58]. Figure 5c and 5d show the variant impact RBPNet profile for SNVs with the lowest associated p-value assigned by Yang et al. for AQR and RBFOX2, respectively. SNVs for both AQR and RBFOX2 are associated with drastic changes in their binding profiles, compared to the references sequences. The sequence around the SNV associated with the most significant binding event of RBFOX2 contains two binding motifs within a 100bp window. When predicting on the reference allele, RBPNet distributes the signal mass equally across the two motifs (5d). As the SNV hits one of the binding motifs, RBPNet in turn predicts a complete flattening of the predicted signal at that motif and a redistribution of the mass to the other motif. This allele-specific effect is clearly displayed both in the predicted profiles and in the attribution maps.

A limitation of Yang et al.’s [58] approach for identifying ASB events is the ambiguity in regards to which of two or more neighboring SNVs is causal for the observed gain or loss of protein-RNA interaction. This is exemplified in Figure **??**, which shows 4 ASB events of QKI within a 200nt interval. As ASB events in [58] are defined based on differential read counts of the REF and ALT allele, reads spanning two or more co-occurring SNVs are counted towards all SNVs, thus leading to “passenger” SNVs that may not be associated with gain or disruption of protein-RNA interactions. Indeed, a separate investigation of impact profiles of each SNVs (Figure **??**) revealed that only 2 (top left / top right) out of 4 SNVs are associated with a substantial change in predicted signal. Interestingly, both SNVs occur in one of the QKI binding motifs visible in the reference attribution maps (Figure **??**). The two remaining SNVs (bottom left / bottom right) fall outside a QKI binding motif and thus show only a negligible change of predicted RBPNet signal. KLD impact scores for SNVs impacting QKI motifs are considerably higher (0.28 and 0.12) compared to SNVs outside of motifs (0.05 and 0.02), which is in line with the strong dependence of allele specific effects of SNVs with respect to their distance to known motifs [58]. Together, these results suggest that variant impact analysis via RBPNet can aid in prioritizing causal SNVs for allele specific binding.

### RBPNet discriminates between functional and non-functional mutations nearby splice junction

Splicing is a complex and tightly regulated post-transcriptional processes in higher eukaryotes that requires concerted binding of multiple RBPs via spliceosomal complexes, with sequence variants disrupting regulatory binding motifs being associated with severe deleterious effects [11]. To evaluate RBPNet’s ability to score the impact of sequence variants on RNA-binding of splicing factors, we obtained a set of 232 splicing-associated mutations from MutSpliceDB [40]. Indeed, we observed greater impact scores for splicing mutations compared to their local controls for 22 out of 40 splicing-related RBPs (Supplementary Figure 3a, Methods 5.12.1). Notably, the set of significant RBPs showed an over-representation of spliceosomal RBPs (15 out of 22), concordant with their higher susceptibility to be directly impacted by splicing mutations at splice junctions. Applying the same procedure with DeepRiPe models, only 4 models (out of 30) showed a significant difference in impact scores allowing discrimination between splicing mutations and local negative controls. This is illustrated by the distribution of scores at various relative distance from splicing junctions, as seen for example with the pre-mRNA processing factor 8 (PRPF8, Supplementary Figure 3b), a core protein of the spliceosome. While the DeepRiPe model showed the most significant discrimination between functional and non-functional mutations (adjusted P-value = 2.23 × 10^−3^), we can see that the score distribution from non-functional mutations at very short distance (<10nt) is slightly more elevated than for mutations beyond. On the other hand, the RBPNet models shows a very clear discrimination even at such short distance. We confirmed the improvement of RBPNet over DeepRiPe for scoring variants by comparing the area under the Receiver Operating Curves from each pair of models (Supplementary Figure 3c), showing that RBPNet was a better classifier for 26 of the 30 models in common.

### Leveraging RBPNet to infer human RBP binding on viral RNAs

We lastly evaluate RBPNet’s ability to infer missing eCLIP signal shape on foreign RNAs. Viruses, including SARS-CoV-2, extensively interact with the host’s RBPome [39, 12, 9]. As the large-scale experimental identification of human RBP binding on viral RNA is associated with significant monetary and labor costs, computational imputation of binding sites represents an attractive alternative to identify crucial host factors involved in the virus life cycle. Recently, Labeau et al. [33] experimentally identified binding of QKI to SARS-CoV-2 and reported that QKI-knockout cells are less permissive than wild-type cells.

Here, we utilize RBPNet models trained on eCLIP datasets to predicting binding profiles of human RBPs to SARS-CoV-2 at single nucleotide resolution. Figure 6b shows RBPNet target track predictions for QKI, showing several high-scoring regions across the SARS-CoV-2 sequence. IG attribution maps at sequence windows around the top 5 highest prediction scores revealed the presence of the QKI binding motif, suggesting that the RBPNet-QKI model predicts bona fide binding profiles. To further validate that RBPNet predictions on SARS-CoV-2, we obtained eCLIP data for CNBP from Schmidt et al. [47]. Indeed, RBPNet target track predictions correlate significantly with the control-normalized eCLIP signal (Figure 6a), with an PCC of 0.210. Together, this indicates that RBPNet trained on human eCLIP experiments may be used to extrapolate eCLIP signal shapes to non-human RNA sequences.

**Figure 6:**
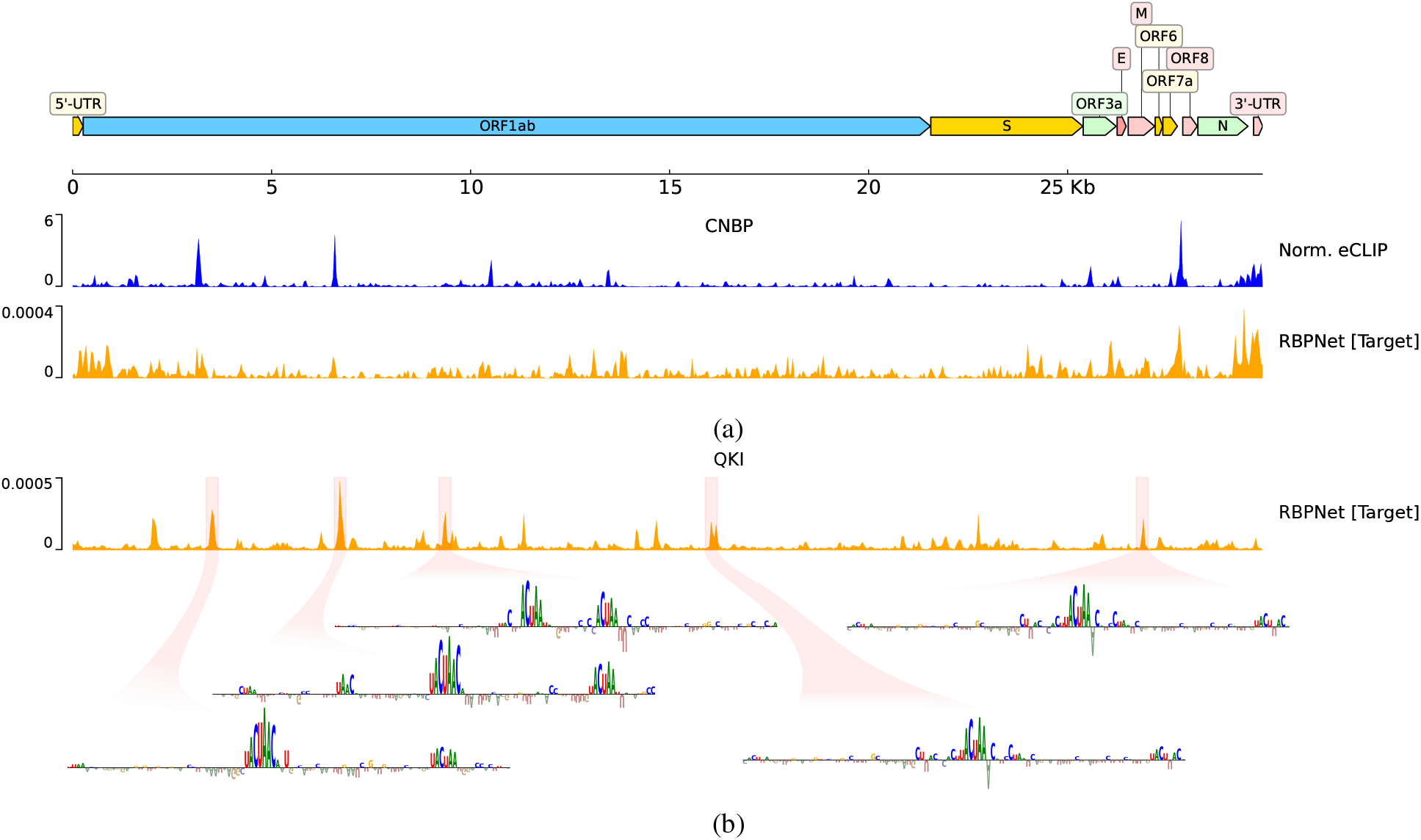
Predicting SARS-CoV-2 binding with RBPNet. The top shows target track predictions using a model trained on CNBP eCLIP data, together with the normalized observed signal taken from Schmidt et al. [47]. Shown below are predictions from a RBPNet-QKI model trained on ENCODE eCLIP data, as well as IG attribution maps around the top-5 positions with highest predicted probability.

### RBPNet Webserver

For ease of use, a *BioLib* webserver version of RBPNet is accessible at https://biolib.com/mhorlacher/RBPNet, enabling prediction of signal profiles and sequence attribution maps with pre-trained models used in this study on user-provided RNA sequences.

## 3 Discussion

While high-throughput single-nucleotide CLIP-seq methods offer unprecedented insights into the protein-RNA interaction landscape of RNA-binding proteins, they are limited to transcripts expressed in the experimental cell type at the time of the experiment. Therefore, researchers must rely on computational methods to impute missing binding information on unexpressed transcripts and or foreign RNAs from sequence. In recent years, an abundance of machine learning methods have been developed for the prediction of protein-RNA interaction from RNA sequence, with the most recent iteration of methods relying on deep neural networks to achieve high state-of-the-art performance. However, current classification-based methods assign predicted binding probabilities to the entire input sequence, typically in the order of hundreds of nucleotides, creating ambiguity with respect to the exact location of protein-RNA interaction. Further, the binary labels used in context of classifiers may only contain a fraction of biological information generated by CLIP experiments, which may hinder the learning of complex associations between RNA sequence and RBP-binding.

The implicit common goal of protein-RNA prediction methods is a near perfect recapitulation of binding information offered by their *in vivo* experimental counterparts, which generate nucleotide-wise counts signal across the transcriptome. In this study, we presented a significant milestone towards this goal in the form of a novel deep learning model, RBPNet, which predicts the CLIP-seq signal shape from RNA sequence at single-nucleotide resolution. By training and evaluating RBPNet on eCLIP, miCLIP and iCLIP datasets, we demonstrated that the model is able to predict the experimental signal shape at high accuracy, reaching replicate-level performance and is applicable to data from different CLIP-based protocols. We leveraged RBPNet’s ability to processes sequences of arbitrary length at prediction time and impute signal shape on entire gene sequences, some of hundreds of kilo-bases in size, and showed that high-scoring positions predicted by RBPNet coincided with single-nucleotide peaks identified from experimental data via the PureCLIP peak caller. Due to their fundamentally different outputs, establishing unbiased comparisons between RBPNet and classification-based models is challenging, as the later assigns predictions to sequence windows rather than individual positions by design. To enable comparison with DeepRiPe, a state-of-the-art CNN classifier, we there generated pseudo position-wise scores via a sliding window approach, and showed that RBPNet significantly outperforms DeepRiPe at recapitation of PureCLIP crosslink sites. This demonstrates that identification of protein-RNA interactions at nucleotide-resolution can not be solved by current classification-based models, stressing the need for new approaches and highlights the novelty of RBPNet, as it is the first method that attempts to prediction protein-RNA interaction at nucleotide resolution.

Unspecific background signal, as well as experimental biases, are an inherent issue of CLIP-based protocols and thus downstream analysis, as bias towards certain sequence elements confounds the learning of genuine, protein-specific sequence features by modeling approaches. For classification-based models, few strategies have been developed to prevent models from learning bias instead of true signal. For instance, Pysster [5] compiles its negative labeled training set for a given RBP by sampling sequences from binding regions of other RBPs, thus explicitly introducing the same sequence biases of the positive set into the negative set, rendering them non-discriminative for the two classes. Similarly, DeepRiPe performs multi-task learning for several RBPs and only trains on sequences that harbor at least one experimentally identified binding site, such that biases associated with CLIP peaks are present in all input sequences. Yan et al. [57] address the RNase T1 cleavage bias towards guanine in the context of PAR-CLIP and HITS-CLIP by replacing nucleotides in a short window up-and downstream of the input sequence viewpoint with uniformly drawn random nucleotides. In contrast to the above strategies, which exclusively rely on manual manipulation of the training data, RBPNet accounts for experimental bias directly as part of its architecture by modeling the control signal as an auxiliary task. Specifically, RBPNet learns a component which explains the difference in signal shape of the experiment and a paired control, the target track, which is expected to be depleted of experimental bias and instead enriched in protein-specific signal. As the *target* signal is not observed experimentally, we instead quantitatively evaluated the ability of RBPNet tracks to recover known RBP binding motifs and showed that the *target* track recovers motifs significantly better than the *total* track, with the later modeling the (possibly biased) observed signal. Orthogonal comparison with 5-mers derived from *in vitro* experiments further showed that the RBPNet *target* track performs comparable to PEKA, a state-of-the-art *de novo* motif discovery tool. This result is remarkable, as RBPNet is not a motif finder by design and the extracted motifs are derived only in retrospect via model interrogation. Both the RBPNet *target* and *total* tracks outperformed DeepRiPe on the task of *in vitro* motif recovery, which highlighted the advantage of learning directly on the bulk of raw experimental signal rather than a compressed representation in the form of binary labels, which may be associated with loss of information. The fact that RBPNet is trained on up to a million (in case of HNRNPC) regions enriched in CLIP-seq signal per RBP may further contribute towards its generalization power. In contrast, classifiers such as DeepRiPe [14] or Pysster [5] are trained on datasets the size of 10-100 thousand samples.

We demonstrated that RBPNet models can be utilized to score the impact of sequence variants on protein-RNA interaction via *in silico* mutagenesis. For instance, we showed that RBPNet scores single-nucleotide variant within known binding motifs or splice junctions significantly higher than randomly selected background variants. This indicates that RBPNet models learned to associated individual nucleotides with the presence or absence of CLIP-seq peaks and that *in silico* probing of the RBPNet model can improve our mechanistic understanding of protein-RNA interaction.

Although we demonstrated the power of our approach on several tasks, including CLIP signal shape imputation, identification of bona fide RBP binding motifs and variant impact scoring, we believe that our results will pave the way for further downstream applications. A promising future application of RBPNet may be *in silico* peak calling for the translation of the predicted signal shape into binding sites -a task that is usually performed by peak calling algorithms in the context of experimental CLIP signal. To achieve that, an open task may be the prediction of absolute signal, i.e. the number of counts falling onto an input sequence rather than their distribution. While RBPNet was shown to predict the counts distribution across an input sequence exceptionally well, the lack of a notion of absolute signal hinders a direct comparison of position-wise probabilities between different RNA sequences. In other words, as RBPNet does not predict the absolute coverage of the CLIP signal, position-wise probabilities are not directly interpretable as RBP-RNA binding strength. On the other hand, this issue is not exclusive to RBPNet, with CLIP peak calling algorithms suffering from a similar issue due to observed position-wise crosslink counts being subject to transcript abundances. Therefore, many peak callers compute position-wise binding site thresholds on a transcript-to-transcript basis (Clipper^4^) or with respect to the local neighborhood (Clippy^5^, Paraclu^6^), a strategy that may be adapted for RBPNet. Nevertheless, future work may explore the feasibility of absolute signal prediction, for instance by incorporating transcript abundance coefficients as model covariates. Avsec et al. [2] demonstrate that the BPNet model may be used to unravel the motif syntax underlying TF cooperativity. In a similar way we envision, as future direction, the application of RBPNet to discover the sequence rules of RBP cooperativity, something that has been only rarely addressed by previous studies. However, CLIP datasets vary greatly in quality, with respect to sequencing depth, number of replicates, replicate consistency, signal-to-noise ratio, and presence or absence of control libraries. While the unique feature of RBPNet to disentangle true signal from noise can in principle enable the accurate identification of composite binding motifs and RBP cooperativity while mitigating the effect of confounding sequence bias, a systematic evaluation of CLIP-seq dataset quality will be necessary to achieve this goal.

## 4 Conclusion

We presented RBPNet, a sequence-to-signal model that predict the distribution of crosslinking events across an input RNA sequence at single-nucleotide resolution. Training and evaluation of RBPNet on 103 eCLIP datasets showed high performance of RBPNet in terms of signal shape correlation, while evaluation on miCLIP and iCLIP datasets demonstrated the models generalization to other CLIP-based protocols. We utilized RBPNet’s ability to handle variable-length input sequence to perform inference on whole-transcript and showed that predicted high-probability positions coincide with PureCLIP peaks, outperforming state-of-the-art classifiers. To account for experimental biases, we additionally modeled the signal distribution of paired control experiments and derived a de-biased component, the RBPNet *target* track, which is enriched in protein-specific signal. We showed that feature importance analysis of the de-biased RBPNet *target* yielded informative sub-sequences which recall *in vitro* motifs at levels comparable to state-of-the-art motif detectors. Finally, we demonstrated RBPNet’s ability to score the impact of SNVs on protein-RNA interaction, which enables prioritization disease-associated variants that disrupt regulatory RNA sequence by causing gain or disruption of RBP binding sites. RBPNet represents a significant milestone towards full *in silico* imputation of protein-RNA interaction, while model interpretation suggest that learning on the raw CLIP signal captures more experimental variants, improving our mechanistic understand of protein-RNA interaction.

## 5 Methods

### 5.1 Data and Preprocessing

#### ENCODE eCLIP

A total of 103 enhanced CLIP (eCLIP) datasets across 103 RBPs from the HepG2 cell line, were obtained from the ENCODE database [[53]]. Each dataset consists of an eCLIP experiment with two replicates and one size-matched input (SMInput) control experiment, which omits the protein-specific immunoprecipitation step and is thus enriched in unspecific background signal. For each eCLIP and SMInput experiment, aligned R2 reads, (i.e. reads who’s start positions likely correspond to the position immediately downs-stream of the RBP cross-linking site) were extracted from the experiment BAM file via SAMtools [[35]]. Next, reads obtained from the both eCLIP replicates were merged and 5’ read-start coverage for both plus and minus strands was computed via BEDtools [[44]].

#### miCLIP

*m*^6^*A* individual-nucleotide resolution UV crosslinking and immunoprecipitation (miCLIP) datasets for HEK293T and mESC cells were obtained from Kortel et al. [30], comprising 4 and 2 replicates, respectively. In addition, miCLIP datasets for the mESC cell line are paired with 2 replicates of a METTL3 KO control experiment. For all datasets, bigWig files of crosslink count signals were directly obtained from the Gene Expression Omnibus (GEO) at the accession number GSE163500. Note that bigWig files of reads without C-to-T transitions were selected, as these reads represent read-through events and would result in unspecific truncation count signal. Lastly, replicates of each dataset were merged by summing of the position-wise crosslink counts.

#### iCLIP

Individual-nucleotide CLIP (iCLIP) datasets for TDP43 and PTBP1, each with two replicates and without control experiments, were obtained from Hallegger et al. [18] and Haberman et al. [15], respectively. Replicates were downloaded from the Sequence Read Archive (SRA) with accession codes ERS10930255 and ERS10930256 for TDP43 and ERR1588764 and ERR1588765 for PTBP1, and processed as described in [56]. The source code for the processing pipeline is available at https://github.com/ulelab/ncawareclip.

### 5.2 Selecting Candidate Sites for Training

In order to speed up convergence of RBPNet, it is important to restrict model training to regions with significant crosslink count signal. In the context of BPNet [2], the authors therefore performed peak calling on ChIP-nexus data to select a set of regions highly enriched in count signal. However, recent work by Toneyan et al. [51] suggests that peak callers select sites too conservatively, which may result in under-fitting of sequence-to-signal models.

To train RBPNet on ENCODE eCLIP datasets, we select a large set of candidate sites as follows. Given the set of genes retrieved from the GENCODE (version 40) [10], a sliding window of size 100 is shifted over each gene (*stride* = 1) and the total number of counts within each window, as well as the highest positional count is obtained. Next, a p-value is computed for each window via a Poisson test by comparing the observed window counts to the expected counts, given the gene-level crosslink counts and the gene-length. At each step, windows with a p-value *<* 0.01, a minimum window count of *N* = 8 and a minimum count *height* (i.e. maximum position-wise count within the window) of *H* = 2 are recorded as candidates and the sliding window is shifted forward by 50 nucleotides. This avoids clusters of redundant candidate sites within transcript regions. Finally, selected 100*nt* windows are extended symmetrically up-and down-stream to a final length of 300*nt*. Note that since RNA is a stranded molecule, counts are obtained for each gene in a strand-specific manner.

For miCLIP datasets, similar parameters together with GENCODE vM23 for the mESC cell line where used. For iCLIP datasets, the minimum window threshold was reduced to *N* = 4, due to a lower sequencing depth.

### 5.3 RBPNet Architecture

The body model architecture of RBPNet was inspired by BPNet [2]. RBPNet takes as input a 300nt RNA sequence, which is one-hot encoded by mapping the bases A, C, G and U to binary vectors [1, 0, 0, 0], [0, 1, 0, 0], [0, 0, 1, 0] and [0, 0, 0, 1], respectively. The 300 × 4 dimensional input is then fed into a 1D convolution layer with 128 filters of size 12, followed by 9 residual blocks. Each residual block consists of (1) a 1D dilated convolution layer with 128 filters of size 6 and exponentially increasing dilation factor, (2) a batch normalization layer, (3) a ReLU activation and (4) a dropout layer with a dropout rate of 0.25, respectively. The output of the last residual block (hereafter referred to as ’bottleneck’ layer) then serves as input to one or more output heads, where each output head corresponds to one of the modeling tasks, for instance the prediction of eCLIP signal and (optionally) SMInput shape. This is outlined schematically in Figure 1b. Each output head consist of a transposed 1D convolutional layer with a single filter of size 20, mapping the bottleneck feature map to a 300-dimensional output vector, which corresponds to the position-wise probabilities of the count distribution within the input window. Notably, as in BPNet [2], same-padding and no pooling is used across all convolution operations in order to conserve the one-to-one correspondence of input sequence positions, feature maps and outputs.

### 5.4 RBPNet Training

Prior to model training, candidate sites obtained in 5.2 were split chromosome-wise into validation (chr2, chr9, chr16), hold-out (chr1, chr8, chr15) and train (all other autosomes) sets for both human and mouse cell lines. RBPNet is trained using the Adam optimizer [[28]] and an initial learning rate (LR) of 0.004. Training is performed for a maximum of 50 epochs with an early-stopping criteria such that training terminates prematurely if the validation loss did not decrease within the last 10 epochs. In addition, a LR schedulers is used such that the LR is halved each time the validation loss did not improve within the last 6 epochs.

### 5.5 RBPNet Loss

The output vector 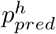 of each track *h* (e.g. eCLIP or SMInput) is used to parameterize a multinomial distribution of read-start counts. For a given training instance, the loss is then computed as the negative log-likelihood of the observed (true) counts 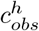, given the total counts 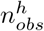 in the input region and the probability vector 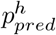. That is, the model’s loss *L*^*h*^ on a task *h* is defined as

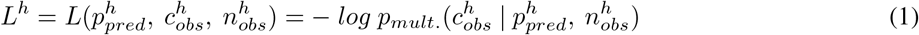

The total loss *L* is then obtained by taking the sum over all task-specific losses.

### 5.6 RBPNet Bias Correction

Experimental bias can lead to unspecific eCLIP signal, severely impacting the downstream binding preference analysis. Therefore, CLIP-seq experiments are usually paired with a control experiment to measure the abundance of background signal at each locus. Assuming that a single read-start count is observed either due to true protein-specific (*target*) signal or experimental bias (*control*), RBPNet models the total CLIP-seq signal as an additive multinomial mixture of the target and bias distributions. That is,

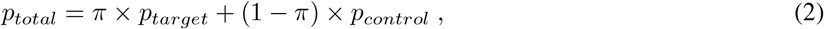

where *p*_*total*_ is the probability vector of the total (e.g. eCLIP) signal, while *p*_*control*_ and *p*_*target*_ are the probability vectors of the control and (unobserved) target signal, respectively. Further, *π/*(1 − *π*) is the relative intensity of the target over the bias signal, given by a mixing coefficient *π*. Note that *π, p*_*target*_ and *p*_*control*_ are learned directly from sequence for each RBP. To ensure that *p*_*control*_ properly approximates the *control* signal distribution over the input sequence, a combined loss on the *total* and *control* tracks is defined as

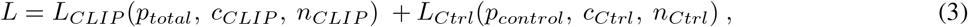

such that *p*_*control*_ is penalized to match the distribution of counts in the control experiment. Once the RBPNet model is trained we can obtain an approximation of the bias-free signal component as *p*_*target*_. A graphical outline of a RBPNet forward pass with bias correction via an additive mixture of *target* and *control* signal is shown in Supplementary Figure 1.

### 5.7 Estimating the Additive Mixing Coefficient

The contribution of target and bias signal towards the total signal is expected to be dependent on the input sequence. For instance, in the presence of multiple RBP binding motifs, the majority of counts may be observed due to protein-specific crosslinking, while under absence of clear binding motifs, crosslinking biases may dominate. RBPNet therefore estimates the multinomial mixing coefficient *π* from the input sequence. Given the feature map of the bottleneck layer (Section 5.3), filter-wise global average pooling is performed along the sequence axis. The resulting 128-dimensional representation of the sequence is then fed into a 1-unit dense layer with linear activation to predict the logit of the mixing coefficient.

### 5.8 Disentangling Target and Control Signals

Given equation 2 together with the total number of counts *N*, we can disentangle the eCLIP signal into the expect control and target counts:

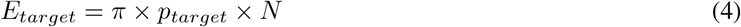

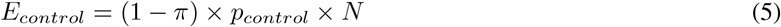

### 5.9 Sequence Importance Scores

To identify RNA sequence features that contributed significantly to the predicted signal distribution, we compute integrated gradients (IG) attribution scores [[50]] of input sequence with respect to the output probability vector *p* for each track. This way, we obtain separate attribution maps for predicted total, target and control signals.

By default, the IG attribution method assumes a classification-based setting, where gradients are computed with respect to the output probability of a target class of interest. For instance in the context of classification-based models, attributions may be computed with respect to a single output neuron describing the binding probability of the target RBP to the input RNA. Here, the resulting feature importance values quantify how much each feature contributed towards the target class. For instance, DeepRiPe employs IG to identify nucleotides that were contributed towards predicting an input sequence as “bound” for a target RBP. In contrast to classification-based methods, RBPNet predicts a 1D profile for each RBP and input sequence. Computing IG attribution maps with respect to only a single position in the output track may draw an incomplete picture of nucleotide-wise contributions towards the predicted signal footprint. We therefore introduce a generalization of IG from scalar to 1D profile outputs.

Given an observed input *x* and a baseline input *x*^′^, the IG score of an input feature *x*_*i*_ is defined as

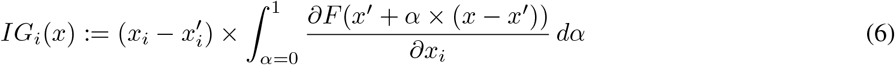

where *F* is a scalar function. In the simplest case of binary classification, where the deep neural network *f* has a scalar output, *F* = *f*. In the case of multi-class classification, *F* is usually defined as *F*(*x*) = Σ_*i*_*p*_*i*_*y*_*i*_, where *p* = *f* (*x*) is the multi-class probability vector and *y* the true label vector with *y*_*i*_ 0, 1, such that IG scores of *x* are obtained with respect to its true class. A natural extension of *F* to count data is given by

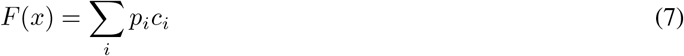

where *p* is the multinomial probability vector and *c* is the vector of true counts, with the desirable effect that predictions at positions with high counts will dominate the input gradients. The extension of *F* in (8) has two major drawbacks. First, it requires true counts *c* for a given sequence to compute attribution scores and second, it might up-weight positions with high counts that are due to experimental bias. We thus reformulate *F* as

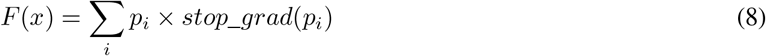

where *stop*_*grad*(*x*) stops gradients flow and treats *x* as a constant.

In other words, instead of computing gradients with respect to the scalar of a single output neuron, we compute gradients of the RNA sequence nucleotides with respect to the sum of the output profile, weighted by a constant version of itself. The weighting ensures that output positions with high probability contribute more towards the nucleotide-wise feature importance scores than low-probability positions.

Note that the proposed generalization of Integrated Gradients to output probability vectors is in analogy to Avsec et al.’s generalization of DeepLIFT [48] scores, described in [2].

By computing gradients with respect to *p*_*target*_ (rather than *p*_*total*_), we explicitly remove contributions of the sequence towards experimental bias and thus focus solely nucleotides that contribute towards the protein-specific crosslinking signal. In general, attribution scores of the total, target and control tracks may be disentangled via

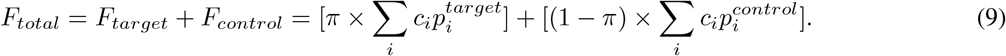

### 5.10 Performance Evaluation

#### 5.10.1 Pearson Correlation Performance

Given the set of 300nt sequences in the hold-out test set, Pearson correlation coefficients (PCC) between RBPNet predictions and the observed crosslink counts, merged between both replicates, were computed. For each eCLIP experiment, the final PCC performance metric is obtained by taking the mean PCC across all test-set sequences.

#### 5.10.2 Comparison with PureCLIP Crosslink Sites

PureCLIP is a single-nucleotide peak caller that identifies significant crosslink (CL) sites by fitting a Hidden Markov Model over the CLIP count signal. To further validate profile predictions made by RBPNet, we investigated whether scores at positions within PureCLIP CL sites are significantly higher than scores outside of CL sites. To this end, we performed PureCLIP CL site ’peak’ calling on all ENCODE eCLIP and miCLIP experiments using default parameters. As suggested by Krakau et al. [31], replicate BAM files were merged to enable the use of signal information across all replicates during peak calling. For ENCODE eCLIP and the mESC miCLIP experiment, PureCLIP was additionally provided BAM files of the control experiment to refine the set of CL sites based on significant enrichment over the control. This was omitted for the HEK miCLIP experiment, as no paired control experiment was available.

For each dataset, transcripts on the hold-out chromosomes (5.4) were intersected with PureCLIP CL sites and only transcripts harboring at least one CL sites were retained, ensuring that the transcript was expressed in the given experiment. Next, whole-transcript predictions were performed with RBPNet, yielding a probability vector of CL enrichment summing up to 1 for each retained transcripts. To measure how well RBPNet predictions discriminate between CL and non-CL sites, the area under the ROC curve (auROC) and the average precision (AP) scores were computed for each transcript. The auROC score may be a more adequate measure of the discriminative power of RBPNet than AP, as the baseline of the AP score is subject to the imbalance of CL and non-CL sites, which are different for each transcript. Further, the auROC is closely related to the Wilcoxon statistic and represents probability of ranking a randomly chosen CL sites above a randomly chosen non-CL site within each transcript, thus directly measuring the discriminative power of the model. Note that within-transcript evaluation is necessary because the position-wise RBPNet scores are subject to transcript length as well as the propensity of RBP binding within the transcript. The finally scalar performance metric for each experiment is then obtained by taking the mean auROC and AP scores across all transcripts.

We additionally evaluated RBPNet against DeepRiPe, a state-of-the-art deep learning model for prediction of protein-RNA interaction, on ENCODE eCLIP datasets. To this end, trained DeepRiPe models for 70 ENCODE HepG2 cell line were obtained from Ghanbari et al. [14]. As DeepRiPe is a classification-based model, the predicted binding probability score is assigned to an entire input regions. To make RBPNet and DeepRiPe comparable on the task of separate PureCLIP CL sites from non-CL sites, we obtained pseudo single-nucleotide resolution scores for DeepRiPe by applying same padding to the transcript sequence, before shifting a sliding window of 150*nt* (DeepRiPe input size) across the sequence and assigning the prediction score to the center position of the current window.

### 5.11 Motif Discovery and Evaluation

#### 5.11.1 RBPmap Motif Evaluation

To quantitatively evaluate the ability of RBPNet to recover known RBP binding motifs in its sequence attribution maps, we compare high-attribution sub-sequences with known binding motifs in the form of position-weight matrices (PWMs) reported in literature. To this end, we first gathered the PWMs of 29 RBPs with both ENCODE eCLIP experiments and reported literature motifs from the RBPmap database [43]. Next, for each eCLIP experiment, the top 5000 ENCODE narrow peaks were selected and profile predictions were performed on a 300*nt* window around the 5’ end of the peak, as this positions has previously been reported to harbor the CL site [8]. After computing attribution maps with respect to the RBPNet *total, target* and *control* tracks, the 5-mer with highest sum of attribution (*IG*_*sum*_) was extracted for each sequence and track. The similarity of each 5-mer to the reference PWM(s) of the RBP was then computed as the mean of the position-wise Jensen–Shannon divergence (JSD) to base 2, a symmetric version of the KL divergence within the bounds [0, 1], where 1 indicates perfect similarity. To account for cases where 5-mers represent truncated motifs or match the reference PWM at a different position offset (i.e. shifted up-or down-stream), we slide each 5-mer over its reference PWM, with a required overlap fraction of 3*nt*. At each shift, the JSD is computed and the final similarity of the 5-mer and the PWM is taken as the maximum similarity over all shift. In cases in which more than one motif PWM is reported for a given RBP in the RBPmap database, similarity computation is performed with respect to all PWMs and the final similarity score is taken as the maximum similarity between the 5-mer and all PWMs of that RBP.

#### 5.11.2 *In Vitro* Motif Evaluation

*In vitro* data on protein-RNA interaction was obtained in the form of k-mer z-scores for RNA-Bind-n-Seq (RBNS) and RNAcompete experiments from Dominquez et al. [8] and Ray et al. [46], respectively. For RBNS, 5mer enrichment scores (*R scores*) for 78 RBPs were obtained from the ENCODE resource, using accession numbers listed in [8]. For each RBP, the *R scores* for the concentration with the highest enrichment were converted to z-scores, by calculating their mean and standard deviation. RNAcompete 7-mer z-scores for 80 RBPs were obtained from Ray et al. [46]. In cases where both RNA-Bind-N-Seq and RNAcompete were available for a particular protein, we prioritized RBNS for downstream analysis, as RBNS z-scores were readily available for 5-mers, whereas RNAcompete required transformation from 7-mer to 5-mer scores. The conversion of 7-mer scores to 5-mer scores was performed by calculating the mean score across all 7-mers that contain a given 5-mer. 7-mers which contain a particular 5-mer more than once were considered as many times as the number of occurrences of the contained 5-mer. For illustration, when calculating the arithmetic mean of z-scores for a 5-mer ‘UUUUU’, the 7-mer ‘UUUUUUG’ would be considered twice (‘[UUUUU]UG’, ‘U[UUUUU]G’). *In vitro* 5-mers of each RBP were then sorted in a descending order based on their enrichment scores and ranked from most (ranked 1st) to least enriched (ranked last).

For evaluation of RBPNet with *in vitro* 5-mers, 5-mers with highest *IG*_*sum*_ were extracted from RBPNet attribution maps of the top 5, 000 ENCODE narrow peaks for each track, as described in 5.11.1. Further, DeepRiPe ENCODE models were obtained from [14] and unique 5-mer counts were obtained in a similar manner by first computing IG attribution maps on a 150*nt* input window around ENCODE narrow peaks and subsequently selecting 5-mers of highest *IG*_*sum*_ for each narrow peak sequence (Methods 5.11.1). For each unique RBPNet and DeepRiPe 5-mer, a relevance score was then computed by taking the sum of *IG*_*sum*_ scores. RBPNet and DeepRiPe 5-mers were then sorted decreasingly with respect to their relevance score. Lastly, we obtained 5-mer enrichment scores calculated with PEKA^7^ (v0.1.6), a motif discovery tool, for all ENCODE eCLIP datasets from [32] (Reference Additional File 5). We used PEKA-scores that were produced with Clippy peaks [56] to rank the k-mers from most to least enriched. As DeepRiPe models were only available for 70 out of 103 ENCODE HepG2 RBPs, and only 27 of those had orthogonal *in vitro* data available, the evaluation of recall was therefore restricted to those proteins. Out of 27 eCLIP datasets, 16 were compared to RBNS and 11 were compared to RNAcompete for recall analysis. Finally, an *in vitro* recall score was computed for each RBP and method by taking the proportion of top 20 5-mers from the corresponding RBNS or RNAcompete dataset that were recovered among the top 20 5-mers in eCLIP, as ranked by the RBPNet tracks, DeepRipe and PEKA.

#### 5.11.3 Consensus Motif Construction

Representative consensus motifs for each eCLIP library were constructed as follows. Given the set of k-mers obtained in Section 5.11.1, k-mers were first sorted by their *IG*_*sum*_ score in descending order. Iterating from the top of the list, the first k-mer is used to seed an initial motif alignment, with consecutive k-mers being aligned (without gaps) to the seed k-mer by sliding the given k-mer over the seed alignment and requiring a minimum overlap of 3*nt*. If no alignment with at least 3 matches is found, the k-mer is considered non-alignable and is instead used to seed a new motif alignment. Consecutive k-mers are aligned to seed alignments in the order of their creation and, if no sufficient alignment is found, are used to seed further motifs alignments on-the-fly. Subsequently, consensus motifs are constructed for each alignment by computing the position-wise nucleotide frequencies within alignments. Consensus motifs can then be prioritized based on the number of supporting k-mers in the underlying alignment. The motif finding procedure was implemented as part of the following Git repository: https://github.com/mhorlacher/metamotif

### 5.12 Variant Impact Scoring

The predicted distribution of counts by RBPNet is solely driven by the input RNA sequence and thus one expects that single-nucleotide variants (SNVs) that fall within crucial sequence feature, such as binding motifs, will have a profound impact on the predicted signal footprint. Therefore, to approximate the impact of a SNV on RBP binding, we quantify the change of the distribution of counts of the alternative allele compared to the reference. To this end, we define the change of count distribution with respect to a given SNV as the KL divergence of the prediction on the SNV-associated allele from the prediction on the reference allele. The variant impact score *KLD*_*SNV*_ is thus defined as

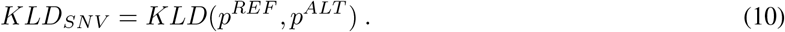

#### 5.12.1 Variant impact scoring of splicing mutations

40 out of 103 RBPs with trained RBPNet-eCLIP models were manually annotated as related to splicing, following the annotations from Nostrand et al. [55] and the HGNC spliceosomal complex groups from the HGNC database [52]. Of these, 21 were further annotated as directly involved in the spliceosome. Next, a set of 260 experimentally validated splicing-related mutations was obtained from MutSpliceDB [40]. After excluding mutations with a distance of more than 10nt from splicing junctions (defined as the first two and last two positions from intronic regions in human coding genes of GENCODE V40 [10]), a set of 232 mutations was retained. For negative controls, we retrieved 6, 087 mutations from gnomAD v2.1.1 ([24]) located within 100nt up-or down-stream of the retained splicing mutations. Control mutations which intersected with the set of splicing-associated mutations were filtered out. Subsequently, RBPNet impact scoring (5.12) was performed on all mutations. For each RBP, a one-sided Wilcoxon ranked sum test was performed to evaluate the enrichment of high-impact splicing-associated mutations over control mutations. P-values were corrected for multiple testing via Benjamini-Hochberg correction, and significance was tested for *α* = 0.05. The same procedure was applied for 30 of the 70 DeepRiPe models found in common with the 40 RBPNet models, taking the models trained from ENCODE HepG2 using both sequence and genomic annotations. Here, the impact score was calculated as the absolute difference in prediction score for a given RBP between the alternative and the reference allele.

## 6 Declarations

### Ethics approval and consent to participate

Not applicable.

### Consent for publication

Not applicable.

### Availability of data and materials

Code for RBPNet training, evaluation, feature importance analysis and variant impact scoring is available at https://github.com/mhorlacher/rbpnet. Code for consensus motif construction is available at https://github.com/mhorlacher/metamotif. Code in both repositories is available under the MIT license.

All data processed in this study was obtained exclusively from public sources. CLIP-seq data was obtained from ENCODE [53] (eCLIP), Kortel et al. [30] (miCLIP) and Hallegger et al. [18] and Haberman et al. [15] (iCLIP). *In vitro* data on protein-RNA interaction was obtained from Dominquez et al. [8] (RNA-Bind-n-Seq) and Ray et al. [46] (RNAcompete). Splicing-related mutations were obtained from MutSpliceDB [40], while further mutations were obtained from gnomAD [24]. Information on allele-specific binding events was taken from Yang et al. [58].

### Authors’ contributions

M.H. and A.M. conceived the project. M.H. collected and processed the datasets, conceptualized and implemented the RBPNet model with help from O.W.; N.W. and L.M. performed analysis of allele-specific binding events and splice-site mutations, respectively, with help from M.H.; K.K. performed binding motif recall analysis on *in vitro* binding data. N.G. helped processing datasets and training RBPNet models. O.W. and A.M. supervised and guided the project, with inputs from M.S., J.U. and J.G.; M.H. wrote the manuscript with help from K.K., N.W., A.M. and L.M. and inputs from O.W., J.G. and J.U.; All authors reviewed and approved the final manuscript.

## Acknowledgements

Not applicable.

## Supplementary Text

### Variance of replicates

Assume that *x* and *y* are repeated measurements with (unknown) mean *m* and independent noise with variance *v*. We have an estimate of the mean 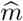. We can calculate the expected squared derivation of the two replicates:

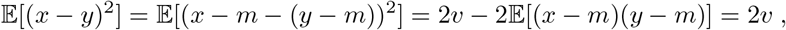

where the cross term disappears because the noise is assumed to be independent on the two replicates. We can also calculate the squared difference between the estimate and a replicate:

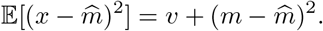

This results follows from the same type of reasoning as above. From these two results we can see that if our estimator is equal to the true mean: 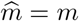 then:

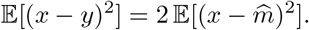

It follows that as the estimated mean 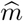 approaches the true mean, the squared derivation of the two replicates exceeds the squared derivation of a replicate and 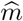 by *v*.

## Supplementary Figures

**Supplementary Figure 1:**
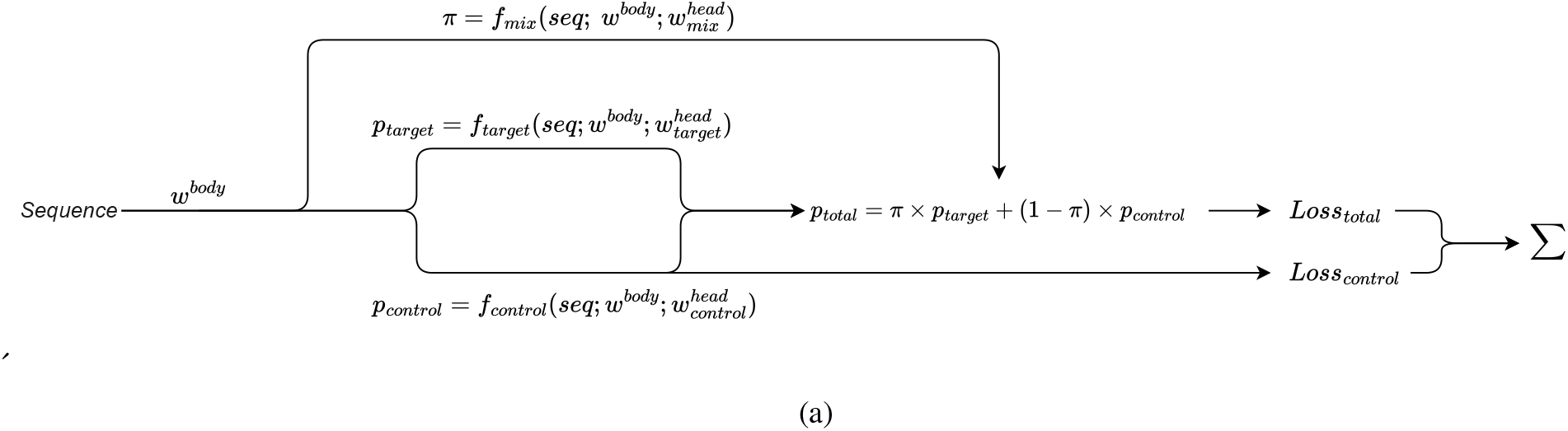
RBPNet forward pass (with bias correction). The feature representation of the input RNA sequence (parameterized by *w*^*body*^) is used to compute the count distribution of the target (*p*_*target*_) and control (*p*_*control*_) tracks. The count distribution of the total track is then given by an additive mixture of target and control tracks, as well a mixing coefficient *π* ∈ [0, 1]. Given the observed counts of the CLIP and control experiment, losses are computed using the total and control tracks, respectively. The total loss is then given by the sum of the total and control track losses.

**Supplementary Figure 2:**
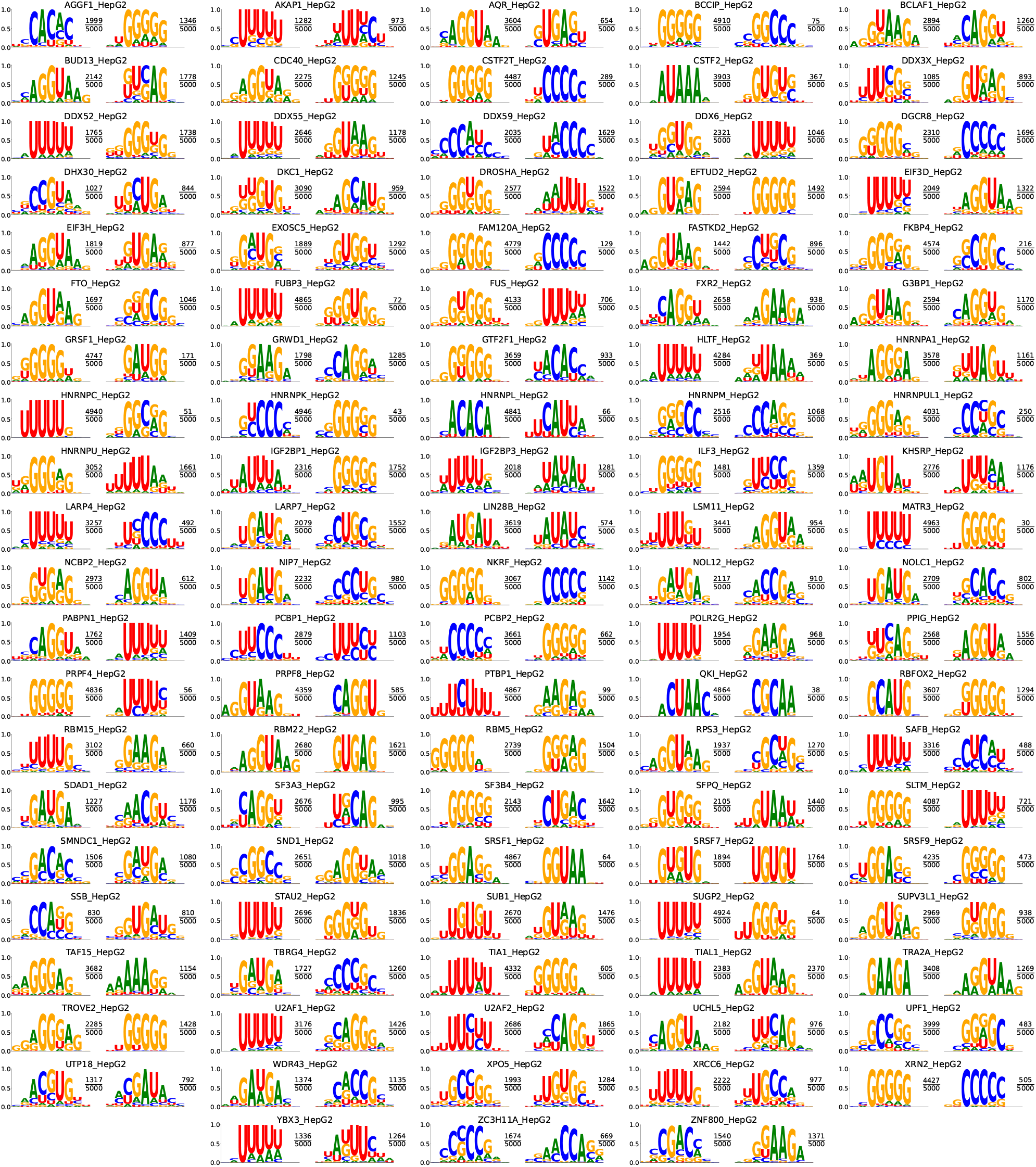
Consensus motifs with highest and second-highest 5-mer support for all 103 ENCODE HepG2 eCLIP experiments.

**Supplementary Figure 3:**
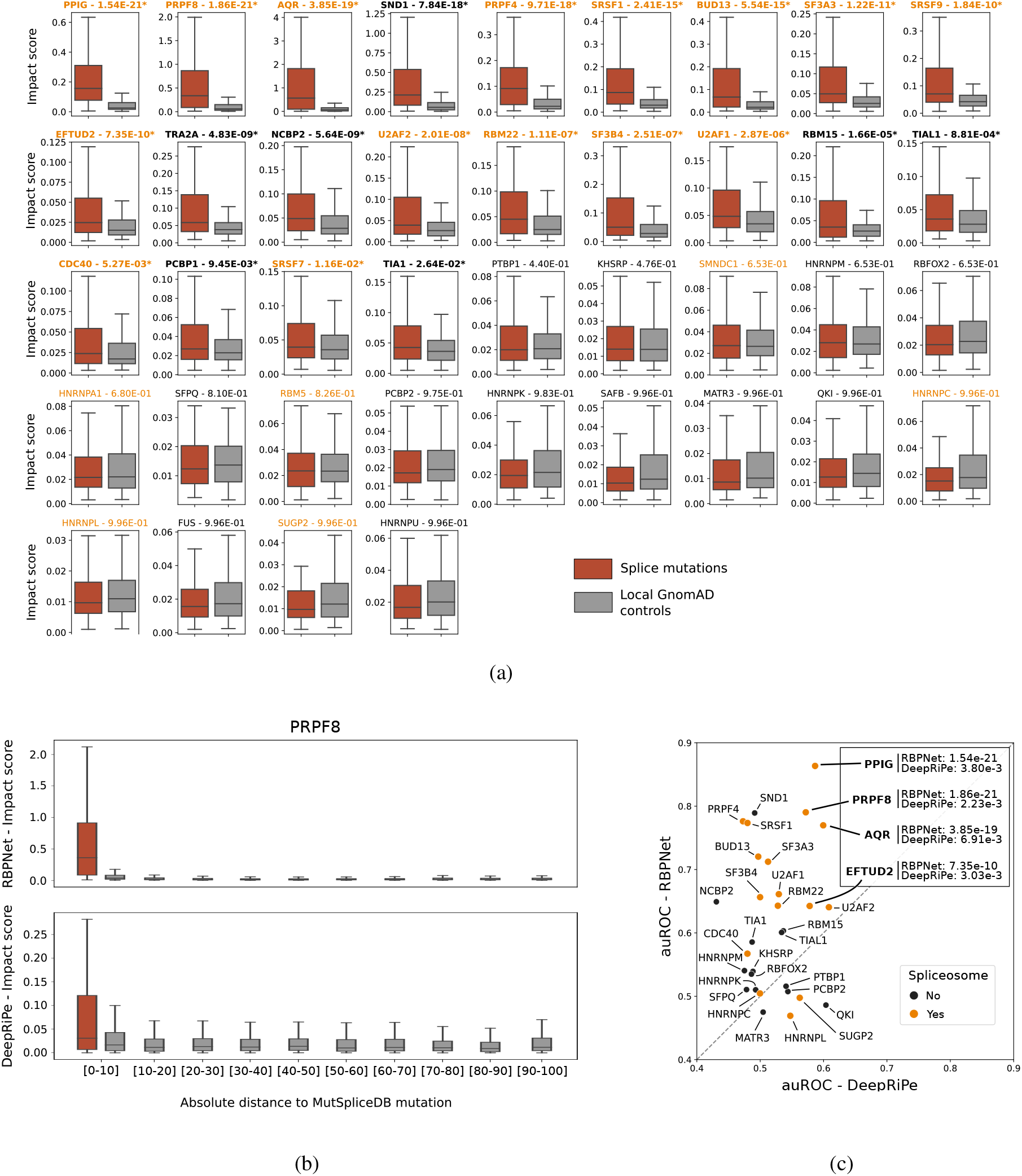
Scoring of 232 splicing mutations from MutSpliceDB along with 6,087 control mutations from gnomAD taken in their vicinity, using 40 splicing-related RBPNet models. **A** Boxplots of RBPNet impact scores from splicing mutations and local controls per RBP. The 40 RBPs are ordered by their adjusted (Benjamini/Hochberg) P-values from Wilcoxon Signed Rank tests. Title in bold: RBPs with significant P-values at *α* = 0.05 (22/40); orange font color indicates the spliceosomal RBPs. **B** Impact score distribution for splice mutations (red boxplot) and gnomAD control mutations (grey boxplots) per absolute, relative-distance bin, with impact scores obtained from RBPNet and DeepRiPe models for the spliceosomal RBP PRPF8 (being the most significant model from DeepRiPe following the Wilcoxon Signed Rank tests). **C** Scatterplot of area under the Receiver Operating Curve (auROC) calculated from RBPNet models (y) and DeepRiPe models (x). All 30 RBPs annotated as splicing-related in common are depicted, with spliceosomal RBPs in orange. Four RBPs are specifically highlighted for being the only models showing significant P-values from the Wilcoxon signed rank tests applied on DeepRiPe models, and their adjusted P-values are reported (along the ajusted P-values for RBPNet.)

## Supplementary Tables

**Supplementary Table 1:** AP and auROC performance of RBPNet (target track) on 103 ENCODE eCLIP experiments from the HepG2 cell line.

|  | RBP_CELL | AP | auROC |  | RBP_CELL | AP | auROC |
| --- | --- | --- | --- | --- | --- | --- | --- |
| 1 | AGGF1_HepG2 | 0.02 | 0.83 | 53 | NKRF_HepG2 | 0.05 | 0.89 |
| 2 | AKAP1_HepG2 | 0.05 | 0.88 | 54 | NOL12_HepG2 | 0.15 | 0.81 |
| 3 | AQR_HepG2 | 0.15 | 0.83 | 55 | NOLC1_HepG2 | 0.18 | 0.76 |
| 4 | BCCIP_HepG2 | 0.08 | 0.95 | 56 | PABPN1_HepG2 | 0.05 | 0.89 |
| 5 | BCLAF1_HepG2 | 0.10 | 0.94 | 57 | PCBP1_HepG2 | 0.09 | 0.87 |
| 6 | BUD13_HepG2 | 0.09 | 0.92 | 58 | PCBP2_HepG2 | 0.14 | 0.97 |
| 7 | CDC40_HepG2 | 0.05 | 0.87 | 59 | POLR2G_HepG2 | 0.09 | 0.97 |
| 8 | CSTF2_HepG2 | 0.12 | 0.93 | 60 | PPIG_HepG2 | 0.11 | 0.96 |
| 9 | CSTF2T_HepG2 | 0.07 | 0.94 | 61 | PRPF4_HepG2 | 0.11 | 0.95 |
| 10 | DDX3X_HepG2 | 0.09 | 0.95 | 62 | PRPF8_HepG2 | 0.22 | 0.97 |
| 11 | DDX52_HepG2 | 0.09 | 0.87 | 63 | PTBP1_HepG2 | 0.25 | 0.93 |
| 12 | DDX55_HepG2 | 0.07 | 0.89 | 64 | QKI_HepG2 | 0.17 | 0.96 |
| 13 | DDX59_HepG2 | 0.05 | 0.87 | 65 | RBFOX2_HepG2 | 0.11 | 0.96 |
| 14 | DDX6_HepG2 | 0.00 | 0.89 | 66 | RBM15_HepG2 | 0.08 | 0.91 |
| 15 | DGCR8_HepG2 | 0.08 | 0.82 | 67 | RBM22_HepG2 | 0.10 | 0.93 |
| 16 | DHX30_HepG2 | 0.04 | 0.87 | 68 | RBM5_HepG2 | 0.07 | 0.84 |
| 17 | DKC1_HepG2 | 0.11 | 0.74 | 69 | RPS3_HepG2 | 0.12 | 0.95 |
| 18 | DROSHA_HepG2 | 0.04 | 0.86 | 70 | SAFB_HepG2 | 0.09 | 0.90 |
| 19 | EFTUD2_HepG2 | 0.04 | 0.84 | 71 | SDAD1_HepG2 | 0.08 | 0.68 |
| 20 | EIF3D_HepG2 | 0.05 | 0.91 | 72 | SF3A3_HepG2 | 0.12 | 0.91 |
| 21 | EIF3H_HepG2 | 0.04 | 0.92 | 73 | SF3B4_HepG2 | 0.12 | 0.96 |
| 22 | EXOSC5_HepG2 | 0.01 | 0.64 | 74 | SFPQ_HepG2 | 0.04 | 0.88 |
| 23 | FAM120A_HepG2 | 0.08 | 0.94 | 75 | SLTM_HepG2 | 0.05 | 0.92 |
| 24 | FASTKD2_HepG2 | 0.06 | 0.90 | 76 | SMNDC1_HepG2 | 0.19 | 0.82 |
| 25 | FKBP4_HepG2 | 0.17 | 0.92 | 77 | SND1_HepG2 | 0.12 | 0.96 |
| 26 | FTO_HepG2 | 0.04 | 0.83 | 78 | SRSF1_HepG2 | 0.12 | 0.97 |
| 27 | FUBP3_HepG2 | 0.10 | 0.93 | 79 | SRSF7_HepG2 | 0.10 | 0.93 |
| 28 | FUS_HepG2 | 0.07 | 0.90 | 80 | SRSF9_HepG2 | 0.15 | 0.94 |
| 29 | FXR2_HepG2 | 0.05 | 0.96 | 81 | SSB_HepG2 | 0.01 | 0.58 |
| 30 | G3BP1_HepG2 | 0.03 | 0.90 | 82 | STAU2_HepG2 | 0.07 | 0.91 |
| 31 | GRSF1_HepG2 | 0.02 | 0.79 | 83 | SUB1_HepG2 | 0.04 | 0.88 |
| 32 | GRWD1_HepG2 | 0.10 | 0.95 | 84 | SUGP2_HepG2 | 0.06 | 0.91 |
| 33 | GTF2F1_HepG2 | 0.10 | 0.90 | 85 | SUPV3L1_HepG2 | 0.09 | 0.74 |
| 34 | HLTF_HepG2 | 0.04 | 0.89 | 86 | TAF15_HepG2 | 0.07 | 0.89 |
| 35 | HNRNPA1_HepG2 | 0.11 | 0.93 | 87 | TBRG4_HepG2 | 0.00 | 0.70 |
| 36 | HNRNPC_HepG2 | 0.23 | 0.98 | 88 | TIA1_HepG2 | 0.08 | 0.93 |
| 37 | HNRNPK_HepG2 | 0.16 | 0.98 | 89 | TIAL1_HepG2 | 0.10 | 0.95 |
| 38 | HNRNPL_HepG2 | 0.16 | 0.98 | 90 | TRA2A_HepG2 | 0.12 | 0.77 |
| 39 | HNRNPM_HepG2 | 0.09 | 0.97 | 91 | TROVE2_HepG2 | 0.10 | 0.86 |
| 40 | HNRNPU_HepG2 | 0.05 | 0.90 | 92 | U2AF1_HepG2 | 0.12 | 0.95 |
| 41 | HNRNPUL1_HepG2 | 0.05 | 0.83 | 93 | U2AF2_HepG2 | 0.16 | 0.98 |
| 42 | IGF2BP1_HepG2 | 0.07 | 0.94 | 94 | UCHL5_HepG2 | 0.09 | 0.93 |
| 43 | IGF2BP3_HepG2 | 0.04 | 0.93 | 95 | UPF1_HepG2 | 0.04 | 0.89 |
| 44 | ILF3_HepG2 | 0.04 | 0.91 | 96 | UTP18_HepG2 | 0.08 | 0.74 |
| 45 | KHSRP_HepG2 | 0.07 | 0.89 | 97 | WDR43_HepG2 | 0.09 | 0.74 |
| 46 | LARP4_HepG2 | 0.07 | 0.88 | 98 | XPO5_HepG2 | 0.01 | 0.82 |
| 47 | LARP7_HepG2 | 0.20 | 0.79 | 99 | XRCC6_HepG2 | 0.04 | 0.88 |
| 48 | LIN28B_HepG2 | 0.04 | 0.91 | 100 | XRN2_HepG2 | 0.09 | 0.92 |
| 49 | LSM11_HepG2 | 0.04 | 0.87 | 101 | YBX3_HepG2 | 0.02 | 0.82 |
| 50 | MATR3_HepG2 | 0.08 | 0.97 | 102 | ZC3H11A_HepG2 | 0.06 | 0.88 |
| 51 | NCBP2_HepG2 | 0.09 | 0.91 | 103 | ZNF800_HepG2 | 0.09 | 0.95 |
| 52 | NIP7_HepG2 | 0.09 | 0.84 |  |  |  |  |

## Footnotes

2 Note that the QKI-specific signal is partially captured in the bias track. While the SMInput experiment omits the immunoprecipi-tation, subsequent size-selection for the target protein still leads to a modest enrichment of protein-specific crosslink signal

3 Note that not all technical CLIP bias is manifested in the RNA sequence and thus learnable by the RBPNet model. Therefore, only the sequence-component of the bias will be captured in control track IG maps.

4 https://github.com/YeoLab/clipper

5 https://github.com/ulelab/clippy

6 https://gitlab.com/mcfrith/paraclu

7 https://github.com/ulelab/peka

## References

[1] B. Alipanahi, A. Delong, M. T. Weirauch, and B. J. Frey. Predicting the sequence specificities of dna-and rna-binding proteins by deep learning. Nature biotechnology, 33(8):831–838, 2015.

[2] Ž. Avsec, M. Weilert, A. Shrikumar, S. Krueger, A. Alexandari, K. Dalal, R. Fropf, C. McAnany, J. Gagneur, A. Kundaje, et al. Base-resolution models of transcription-factor binding reveal soft motif syntax. Nature Genetics, 53(3):354–366, 2021.

[3] T. L. Bailey, M. Boden, F. A. Buske, M. Frith, C. E. Grant, L. Clementi, J. Ren, W. W. Li, and W. S. Noble. Meme suite: tools for motif discovery and searching. Nucleic acids research, 37(suppl_2):W202–W208, 2009.

[4] S. Bergstrand, E. M. O’Brien, C. Coucoravas, D. Hrossova, D. Peirasmaki, S. Schmidli, S. Dhanjal, C. Pederiva, L. Siggens, O. Mortusewicz, et al. Small cajal body-associated rna 2 (scarna2) regulates dna repair pathway choice by inhibiting dna-pk. Nature communications, 13(1):1–18, 2022.

[5] S. Budach and A. Marsico. Pysster: classification of biological sequences by learning sequence and structure motifs with convolutional neural networks. Bioinformatics, 34(17):3035–3037, 2018.

[6] X. Chen, Y. Liu, C. Xu, L. Ba, Z. Liu, X. Li, J. Huang, E. Simpson, H. Gao, D. Cao, et al. Qki is a critical pre-mrna alternative splicing regulator of cardiac myofibrillogenesis and contractile function. Nature communications, 12(1):1–18, 2021.

[7] L. De Conti, M. Baralle, and E. Buratti. Neurodegeneration and rna-binding proteins. Wiley Interdisciplinary Reviews: RNA, 8(2):e1394, 2017.

[8] D. Dominguez, P. Freese, M. S. Alexis, A. Su, M. Hochman, T. Palden, C. Bazile, N. J. Lambert, E. L. Van Nostrand, G. A. Pratt, et al. Sequence, structure, and context preferences of human rna binding proteins. Molecular cell, 70(5):854–867, 2018.

[9] R. A. Flynn, J. A. Belk, Y. Qi, Y. Yasumoto, J. Wei, M. M. Alfajaro, Q. Shi, M. R. Mumbach, A. Limaye, P. C. DeWeirdt, et al. Discovery and functional interrogation of sars-cov-2 rna-host protein interactions. Cell, 184(9):2394–2411, 2021.

[10] A. Frankish, M. Diekhans, I. Jungreis, J. Lagarde, J. E. Loveland, J. M. Mudge, C. Sisu, J. C. Wright, J. Armstrong, I. Barnes, et al. Gencode 2021. Nucleic acids research, 49(D1):D916–D923, 2021.

[11] A. M. Fredericks, K. J. Cygan, B. A. Brown, and W. G. Fairbrother. RNA-Binding Proteins: Splicing Factors and Disease. Biomolecules, 5(2):893–909, May 2015.

[12] M. Garcia-Moreno, A. I. Järvelin, and A. Castello. Unconventional rna-binding proteins step into the virus–host battlefront. Wiley Interdisciplinary Reviews: RNA, 9(6):e1498, 2018.

[13] F. Gebauer, T. Schwarzl, J. Valcárcel, and M. W. Hentze. Rna-binding proteins in human genetic disease. Nature Reviews Genetics, 22(3):185–198, 2021.

[14] M. Ghanbari and U. Ohler. Deep neural networks for interpreting rna-binding protein target preferences. Genome research, 30(2):214–226, 2020.

[15] N. Haberman, I. Huppertz, J. Attig, J. König, Z. Wang, C. Hauer, M. W. Hentze, A. E. Kulozik, H. Le Hir, T. Curk, et al. Insights into the design and interpretation of iclip experiments. Genome biology, 18(1):1–21, 2017.

[16] M. Hafner, M. Katsantoni, T. Köster, J. Marks, J. Mukherjee, D. Staiger, J. Ule, and M. Zavolan. Clip and complementary methods. Nature Reviews Methods Primers, 1(1):1–23, 2021.

[17] M. Hafner, M. Landthaler, L. Burger, M. Khorshid, J. Hausser, P. Berninger, A. Rothballer, M. Ascano Jr, A.-C. Jungkamp, M. Munschauer, et al. Transcriptome-wide identification of rna-binding protein and microrna target sites by par-clip. Cell, 141(1):129–141, 2010.

[18] M. Hallegger, A. M. Chakrabarti, F. C. Lee, B. L. Lee, A. G. Amalietti, H. M. Odeh, K. E. Copley, J. D. Rubien, B. Portz, K. Kuret, et al. Tdp-43 condensation properties specify its rna-binding and regulatory repertoire. Cell, 184(18):4680–4696, 2021.

[19] D. Heller, R. Krestel, U. Ohler, M. Vingron, and A. Marsico. sshmm: extracting intuitive sequence-structure motifs from high-throughput rna-binding protein data. Nucleic acids research, 2017.

[20] M. W. Hentze, A. Castello, T. Schwarzl, and T. Preiss. A brave new world of rna-binding proteins. Nature reviews Molecular cell biology, 19(5):327–341, 2018.

[21] M. Horlacher, S. Oleshko, Y. Hu, M. Ghanbari, E. E. Vergara, N. Mueller, U. Ohler, L. Moyon, and A. Marsico. Computational mapping of the human-sars-cov-2 protein-rna interactome. bioRxiv, 2021.

[22] I. Huppertz, J. Attig, A. D’Ambrogio, L. E. Easton, C. R. Sibley, Y. Sugimoto, M. Tajnik, J. König, and J. Ule. iclip: protein–rna interactions at nucleotide resolution. Methods, 65(3):274–287, 2014.

[23] K. Izumikawa, Y. Nobe, H. Ishikawa, Y. Yamauchi, M. Taoka, K. Sato, H. Nakayama, R. J. Simpson, T. Isobe, and N. Takahashi. Tdp-43 regulates site-specific 2-o-methylation of u1 and u2 snrnas via controlling the cajal body localization of a subset of c/d scarnas. Nucleic acids research, 47(5):2487–2505, 2019.

[24] K. J. Karczewski, L. C. Francioli, G. Tiao, B. B. Cummings, J. Alföldi, Q. Wang, R. L. Collins, K. M. Laricchia, A. Ganna, D. P. Birnbaum, et al. The mutational constraint spectrum quantified from variation in 141,456 humans. Nature, 581(7809):434–443, 2020.

[25] H. Kazan, D. Ray, E. T. Chan, T. R. Hughes, and Q. Morris. Rnacontext: a new method for learning the sequence and structure binding preferences of rna-binding proteins. PLoS computational biology, 6(7):e1000832, 2010.

[26] S. Ke, A. Pandya-Jones, Y. Saito, J. J. Fak, C. B. Vågbø, S. Geula, J. H. Hanna, D. L. Black, J. E. Darnell, and R. B. Darnell. m6a mrna modifications are deposited in nascent pre-mrna and are not required for splicing but do specify cytoplasmic turnover. Genes & development, 31(10):990–1006, 2017.

[27] D. R. Kelley, Y. A. Reshef, M. Bileschi, D. Belanger, C. Y. McLean, and J. Snoek. Sequential regulatory activity prediction across chromosomes with convolutional neural networks. Genome research, 28(5):739–750, 2018.

[28] D. P. Kingma and J. Ba. Adam: A method for stochastic optimization. arXiv preprint arXiv:1412.6980, 2014.

[29] J. König, K. Zarnack, G. Rot, T. Curk, M. Kayikci, B. Zupan, D. J. Turner, N. M. Luscombe, and J. Ule. iclip reveals the function of hnrnp particles in splicing at individual nucleotide resolution. Nature structural & molecular biology, 17(7):909–915, 2010.

[30] N. Körtel, C. Rücklé, Y. Zhou, A. Busch, P. Hoch-Kraft, F. R. Sutandy, J. Haase, M. Pradhan, M. Musheev, D. Ostareck, et al. Deep and accurate detection of m6a rna modifications using miclip2 and m6aboost machine learning. Nucleic acids research, 49(16):e92–e92, 2021.

[31] S. Krakau, H. Richard, and A. Marsico. Pureclip: capturing target-specific protein–rna interaction footprints from single-nucleotide clip-seq data. Genome biology, 18(1):1–17, 2017.

[32] K. Kuret, A. G. Amalietti, D. M. Jones, C. Capitanchik, and J. Ule. Positional motif analysis reveals the extent of specificity of protein-rna interactions observed by clip. Genome Biology, 23(1):1–34, 2022.

[33] A. Labeau, L. Fery-Simonian, A. Lefevre-Utile, M. Pourcelot, L. Bonnet-Madin, V. Soumelis, V. Lotteau, P.-O. Vidalain, A. Amara, and L. Meertens. Characterization and functional interrogation of the sars-cov-2 rna interactome. Cell reports, 39(4):110744, 2022.

[34] N. Lambert, A. Robertson, M. Jangi, S. McGeary, P. A. Sharp, and C. B. Burge. Rna bind-n-seq: quantitative assessment of the sequence and structural binding specificity of rna binding proteins. Molecular cell, 54(5):887–900, 2014.

[35] H. Li, B. Handsaker, A. Wysoker, T. Fennell, J. Ruan, N. Homer, G. Marth, G. Abecasis, and R. Durbin. The sequence alignment/map format and samtools. Bioinformatics, 25(16):2078–2079, 2009.

[36] B. Linder, A. V. Grozhik, A. O. Olarerin-George, C. Meydan, C. E. Mason, and S. R. Jaffrey. Single-nucleotide-resolution mapping of m6a and m6am throughout the transcriptome. Nature methods, 12(8):767–772, 2015.

[37] D. Maticzka, S. J. Lange, F. Costa, and R. Backofen. Graphprot: modeling binding preferences of rna-binding proteins. Genome biology, 15(1):1–18, 2014.

[38] K. D. Meyer. Dart-seq: an antibody-free method for global m6a detection. Nature methods, 16(12):1275–1280, 2019.

[39] J. M. Molleston and S. Cherry. Attacked from all sides: Rna decay in antiviral defense. Viruses, 9(1):2, 2017.

[40] A. Palmisano, S. Vural, Y. Zhao, and D. Sonkin. MutSpliceDB: A database of splice sites variants with RNA-seq based evidence on effects on splicing. Human Mutation, 42(4):342–345, 2021. _eprint: https://onlinelibrary.wiley.com/doi/pdf/10.1002/humu.24185.

[41] X. Pan, P. Rijnbeek, J. Yan, and H.-B. Shen. Prediction of rna-protein sequence and structure binding preferences using deep convolutional and recurrent neural networks. BMC genomics, 19(1):1–11, 2018.

[42] C. Y. Park, J. Zhou, A. K. Wong, K. M. Chen, C. L. Theesfeld, R. B. Darnell, and O. G. Troyanskaya. Genome-wide landscape of rna-binding protein target site dysregulation reveals a major impact on psychiatric disorder risk. Nature genetics, 53(2):166–173, 2021.

[43] I. Paz, I. Kosti, M. Ares Jr, M. Cline, and Y. Mandel-Gutfreund. Rbpmap: a web server for mapping binding sites of rna-binding proteins. Nucleic acids research, 42(W1):W361–W367, 2014.

[44] A. R. Quinlan and I. M. Hall. Bedtools: a flexible suite of utilities for comparing genomic features. Bioinformatics, 26(6):841–842, 2010.

[45] D. Ray, H. Kazan, E. T. Chan, L. P. Castillo, S. Chaudhry, S. Talukder, B. J. Blencowe, Q. Morris, and T. R. Hughes. Rapid and systematic analysis of the rna recognition specificities of rna-binding proteins. Nature biotechnology, 27(7):667–670, 2009.

[46] D. Ray, H. Kazan, K. B. Cook, M. T. Weirauch, H. S. Najafabadi, X. Li, S. Gueroussov, M. Albu, H. Zheng, A. Yang, et al. A compendium of rna-binding motifs for decoding gene regulation. Nature, 499(7457):172–177, 2013.

[47] N. Schmidt, C. A. Lareau, H. Keshishian, S. Ganskih, C. Schneider, T. Hennig, R. Melanson, S. Werner, Y. Wei, M. Zimmer, et al. The sars-cov-2 rna–protein interactome in infected human cells. Nature microbiology, 6(3):339–353, 2021.

[48] A. Shrikumar, P. Greenside, and A. Kundaje. Learning important features through propagating activation differences. In International conference on machine learning, pages 3145–3153. PMLR, 2017.

[49] Y. Sugimoto, J. König, S. Hussain, B. Zupan, T. Curk, M. Frye, and J. Ule. Analysis of clip and iclip methods for nucleotide-resolution studies of protein-rna interactions. Genome biology, 13(8):1–13, 2012.

[50] M. Sundararajan, A. Taly, and Q. Yan. Axiomatic attribution for deep networks. In International Conference on Machine Learning, pages 3319–3328. PMLR, 2017.

[51] S. Toneyan, Z. Tang, and P. K. Koo. Evaluating deep learning for predicting epigenomic profiles. BioRxiv, 2022.

[52] S. Tweedie, B. Braschi, K. Gray, T. E. M. Jones, R. Seal, B. Yates, and E. A. Bruford. Genenames.org: the HGNC and VGNC resources in 2021. Nucleic Acids Research, 49(D1):D939–D946, Jan. 2021.

[53] E. L. Van Nostrand, P. Freese, G. A. Pratt, X. Wang, X. Wei, R. Xiao, S. M. Blue, J.-Y. Chen, N. A. Cody, D. Dominguez, et al. A large-scale binding and functional map of human rna-binding proteins. Nature, 583(7818):711–719, 2020.

[54] E. L. Van Nostrand, G. A. Pratt, A. A. Shishkin, C. Gelboin-Burkhart, M. Y. Fang, B. Sundararaman, S. M. Blue, T. B. Nguyen, C. Surka, K. Elkins, et al. Robust transcriptome-wide discovery of rna-binding protein binding sites with enhanced clip (eclip). Nature methods, 13(6):508–514, 2016.

[55] E. L. Van Nostrand, G. A. Pratt, B. A. Yee, E. C. Wheeler, S. M. Blue, J. Mueller, S. S. Park, K. E. Garcia, C. Gelboin-Burkhart, T. B. Nguyen, I. Rabano, R. Stanton, B. Sundararaman, R. Wang, X.-D. Fu, B. R. Graveley, and G. W. Yeo. Principles of RNA processing from analysis of enhanced CLIP maps for 150 RNA binding proteins. Genome Biology, 21(1):90, Apr. 2020.

[56] R. A. Varier, T. Sideri, C. Capitanchik, Z. Manova, E. Calvani, A. Rossi, R. Edupuganti, I. Ensinck, V. W. Chan, H. Patel, et al. m6a reader pho92 is recruited co-transcriptionally and couples translation efficacy to mrna decay to promote meiotic fitness in yeast. bioRxiv, 2022.

[57] Z. Yan, W. L. Hamilton, and M. Blanchette. Graph neural representational learning of rna secondary structures for predicting rna-protein interactions. Bioinformatics, 36(Supplement_1):i276–i284, 2020.

[58] E.-W. Yang, J. H. Bahn, E. Y.-H. Hsiao, B. X. Tan, Y. Sun, T. Fu, B. Zhou, E. L. Van Nostrand, G. A. Pratt, P. Freese, et al. Allele-specific binding of rna-binding proteins reveals functional genetic variants in the rna. Nature communications, 10(1):1–15, 2019.

